# The Series of a Four-node Motif and Coherent Type-1 Feed-forward Loop can Provide Sensitive Detection over Arbitrary Range of Input Signal, thereby Explain Weber’s Law in Higher-Order Sensory Processes, and Compute Logarithm

**DOI:** 10.1101/2020.04.08.032193

**Authors:** Dinkar Wadhwa

**Affiliations:** Department of Systems and Computational Biology University of Hyderabad, Hyderabad - 500046, India

**Author notes:** The author did 90% of the work without an affiliation.

**Keywords:** Sensitive detection, Weber’s law, linear model, bow-tie architecture, numerosity processing, human cone cell, logarithm

## Abstract

Weber’s law states that the ratio of the smallest perceptual change in an input signal and the background signal is constant. The law is observed across the perception of weight, light intensity, and sound intensity and pitch. Conventionally, two models of perception have been proposed to explain Weber’s law, namely the logarithmic and the linear model. Later, another formulation of Weber’s law was proposed which links it with exact adaptation. This paper argues in favour of the linear model, which requires the sensory system to generate linear input-output relationship over several orders of magnitude. To this end, a four-node motif (FNM) is constructed from first principles whose series provides almost linear relationship between input signal and output over arbitrary range of input signal. Mathematical analysis into the origin of this quasi-linear relationship shows that the series of coherent type-1 feed-forward loop (C1-FFL) is able to provide perfectly linear input-output relationship over arbitrary range of input signal. FNM also reproduces the neuronal data of numerosity detection study on monkey. The series of FNM also provides a mechanism for sensitive detection over arbitrary range of input signal when the output has an upper limit. Further, the series of FNM provides a general basis for a class of bow-tie architecture where the number of receptors is much lower than the range of input signal and the “decoded output”. Besides (quasi-)linear input-output relationship, another example of this class of bow-tie architecture that the series of FNM is able to produce is absorption spectra of cone opsins of humans. Further, the series of FNM and C1-FFL, respectively, can compute logarithm over arbitrary range of input signal.

## 1. Introduction

Ernst Weber discovered that the ratio of the smallest noticeable difference in an input signal and the background signal is constant, a phenomenon now known as Weber’s law [1]. Later, Fechner showed that Weber’s law could be explained by assuming that the perception is proportional to the logarithm of the input signal (see the section *Weber*’*s Law*) [2]. Another model which can also account for Weber’s law is the linear model, according to which the output is equal to the input signal plus noise which is proportional to the input signal. In 1973, work of Mesibov et al. on bacterial chemotaxis suggested that *Escherichia coli* exhibits Weber’s law, in that the bacterial response was dependent on the gradient of the logarithmic ligand concentration [3]. A theoretical study of a reduced model of *Escherichia coli*’s chemotaxis machinery showed that exact adaptation is necessary, though not sufficient, for the bacterial chemotaxis to exhibit Weber’s law [4]. Later an experimental and theoretical modelling study using the reduced model demonstrated the presence of Weber’s law in *E. coli*’s chemotaxis. [5].

Thus, inspired from the connection between Weber’s law and exact adaptation in the bacterium, Weber’s law was formulated as an exactly adapting system generating identical maximum amplitude as a response to an input signal if the ratio of the current input signal to the background signal is constant [6,7]. The formulation has been applied to explain modified versions of Weber-Fechner and Stevens laws found in human sensory systems [8]. However, it is doubtful that all higher-order sensory processes which show Weber’s law exhibit scale-invariance with respect to the maximum amplitude. For example, consider the case of weight perception. Suppose the just noticeable difference (JND) for weight perception is 5% of the background, so that a subject holding a weight of 100 gm can feel the change in the weight if at least 5 gm of weight is added. The JND when the subject is holding 1 kg of weight is 50 gm. Although the ratio of the JND to the background signal is constant, the subject is able to sense the background weight and the additional weight in the absolute sense. The subject is not, if at all, just calculating the ratio between the two signals.

It has also been claimed that fold-change detection (FCD) entails Weber’s law [6]. A system capable of providing FCD also shows exact adaptation, in which the response to an input signal returns to the resting value of the system even when the signal persists. That FCD implies exact adaptation can be proven in the following way. Consider a system equilibrated at a stimulus X_0_. Its baseline output is Z_0_. It generates a certain output when a stimulus X_1_ is provided. The ratio X_1_/X_0_ is the fold change, f. If the system does not show exact adaptation, the resting output of the system, equilibrated at stimulus X_1_ (or fX_0_), will not be equal to Z_0_ (scenario I). If the system is capable of providing FCD, it will show identical response when the two stimuli are replaced by kX_0_ and kX_1_, respectively, where k is a scalar. Clearly, the output of the system equilibrated at stimulus kX_0_ should be Z_0_ (scenario II). If k is set equal to f, this leads to a contradiction in a system which shows FCD but does not adapt exactly. That is, when the stimulus is fX_0_, the resting output of the system is Z_0_ in the scenario II, whereas it is not equal to Z_0_ in the scenario I. Therefore, a system which shows FCD will also show exact adaptation.

However, it is doubtful that all higher-order sensory processes which show Weber’s law exhibit exact adaptation. Again considering the case of weight perception, a subject holding a certain amount of weight continues to sense the same amount of input signal; that is, the perception of the weight does not come back to some resting value. The same argument applies also to the perception of sound intensity and brightness.

Weber’s law is also observed in size (length/area/volume) perception, for which two examples are presented now. The first example demonstrates that a difference of 1 unit between two rectangular strips of 5 and 6 units is perceptible to the human eye, whereas the same difference between two rectangular strips of 40 and 41 units is not (Figure 2A). The second example shows that the human eye can easily notice the difference of one disc between collections of 4 and 5 such identical discs at a quick glance, whereas doing so is virtually impossible between collections of 24 ad 25 discs (Figure 2B). Weber’s law is also observed in numerosity processing [9]. As with weight perception, the perception of size and number does not come back to some resting value. Because it is not clear how scale-invariance with respect to the maximum amplitude or exact adaptation could be present in weight perception, size perception, and numerosity processing; arguably, a general mechanism for Weber’s law is needed which does not involve either scale-invariance with respect to the maximum amplitude or exact adaptation. Additionally, such a mechanism should be able to generate linear input-output relationship over several orders of magnitude.

**Figure 1:**
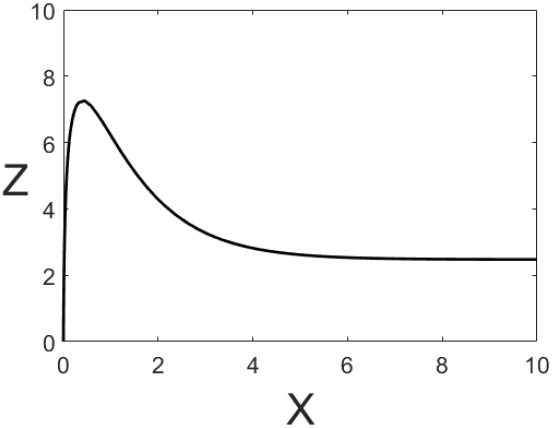
Output of a type-1 incoherent feed-forward loop that lacks exact adaptation.

**Figure 2:**
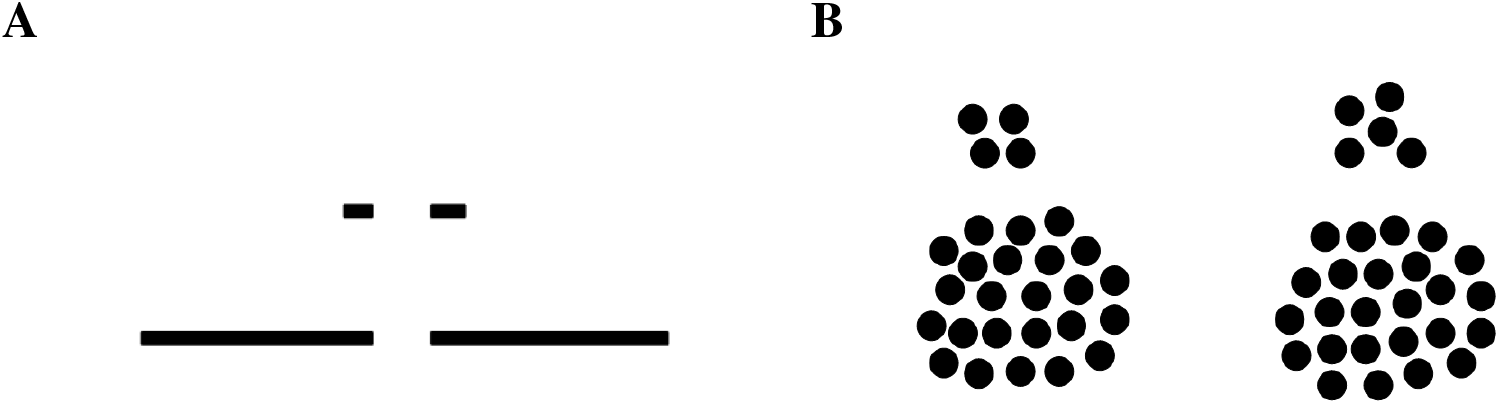
(**A**) Two pairs of rectangular strips which differ by the same length (20% of the smaller strip of the upper pair). (**B**) Two pairs of collection of discs numbering (i) 4 and 5; (ii) 24 and 25.

## 2. Results

### 2.1. (Quasi-)Linear Input-Output Relationship Over Arbitrary Range

In order for a system to respond sensitively throughout the whole range of input signal, ideal relationship between an input signal and the output should be a straight line with the slope equal to 1 and passing through the origin (i.e., the y-intercept equal to zero). Because sensory systems operate over several orders of magnitude, the system should be able to generate linear input-output relationship over several orders of magnitude. This is achieved in the following way. Consider a collection of five receptors which bind a certain ligand, representing the input signal, with the equilibrium dissociation constant (K_da_) equal to 10^1^, 10^2^, 10^3^, 10^4^, and 10^5^ arb. units, respectively. The input signal, X, causes the *Z* producer (an imaginary generic form of gene) to produce Z. In order to have sensitive detection, that receptor should contribute to the output, Z, whose K_da_ is an order of magnitude larger than the strength of the input signal (the concentration of X). Clearly, the output (i.e., the production rate of Z, β_1_) of the receptor meant to respond to an input signal of higher strength should be proportionally larger.

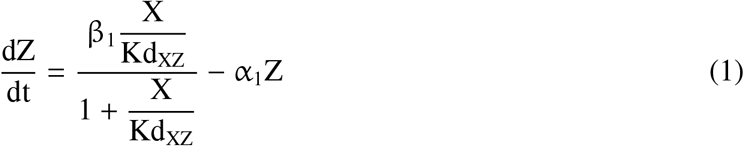

where, β_1_ is the rate constant for production of Z by X. α_1_ is the degradation rate of Z. Kd_XZ_ is the equilibrium dissociation constant for the binding of X to the producer element (an imaginary generic form of promoter) of Z, which causes Z production.

The equilibrium value of Z is

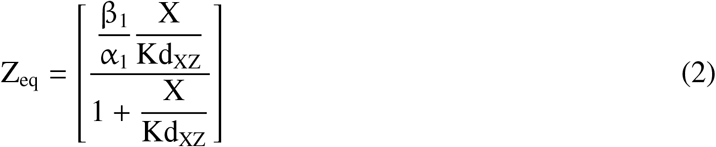

Choosing appropriate values of β_1_/α_1_ and Kd_XZ_, the total output from the N receptors is

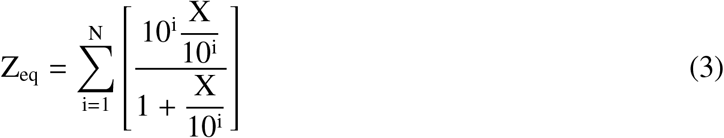

Let’s assume that the input signal’s value has the order of magnitude two. Arguably, the receptor with Kd_XZ_ equal to 10^3^ should generate output from the input signal of this value. However, the receptors with Kd_XZ_ equal to 10^1^ and 10^2^ arb. units, respectively, largely saturated by the input signal, also contribute to the output. Thus, output from the receptors whose Kd_XZ_ is either equal to or less than the order of magnitude of the input signal should be removed. This is achieved by adding to the motif a “negative arm”: the input signal X engenders production of the molecule Y which suppresses the output Z from such receptors. The input signal X and the molecule Y bind to the producer element of Z in mutually exclusive manner. The resulting motif is type-1 incoherent feed-forward loop. Parameters of the negative arm are such that it suppresses the output mainly from the receptors whose Kd_XZ_ is either equal to or less than the order of magnitude of the input signal.

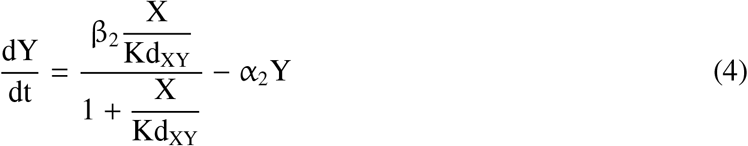

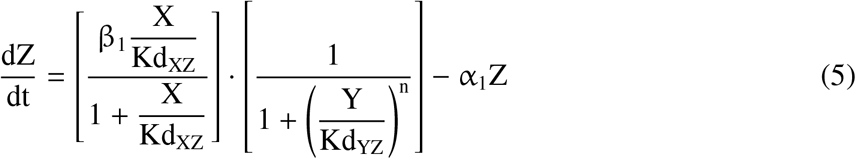

where, β_2_ is the rate constant for production of Y by X. α_2_ is the degradation rate of Y. Kd_XY_ is the equilibrium dissociation constant for the binding of X to the producer element of Y, which causes Y production. Kd_YZ_ is the equilibrium dissociation constant for the binding of Y to the producer element of Z, leading to suppression of Z production.

The equilibrium value of Y and Z are

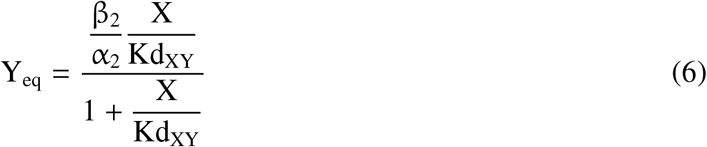

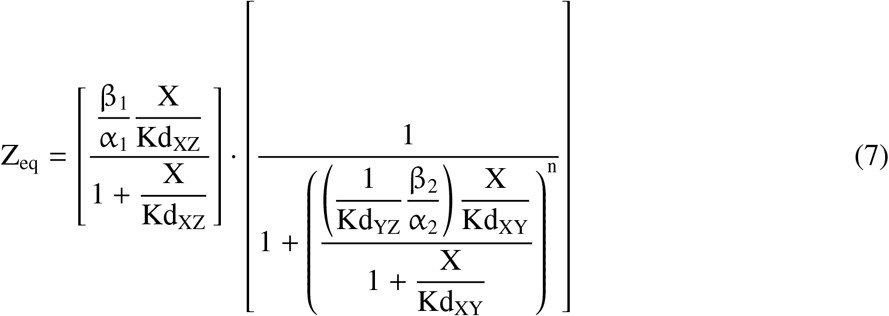

The expression in the second pair of brackets on the right hand side (RHS) of eq. 7, representing suppression of the output by the input signal, reaches an asymptotic value because the denominator in the expression reaches an asymptotic value. On the other hand, the formulation demands the expression to keep decreasing with increasing X. That is, the expression should be of the form: 1/(1 +X^n^). This is achieved in the following way. The expression in the second pair of brackets on the RHS of eq. 7 is approximately equal to the expression 8 (which is of the required form), at least in the parameter range of interest (i.e., until X is not at least one order of magnitude larger than Kd_XZ_), if Kd_XY_ ≫ Kd_XZ_ and the order of magnitude of (1/Kd_YZ_)·(β_2_/α_2_) is at least four (Figure 3A). This is because when Kd_XY_ ≫ Kd_XZ_, X/Kd_XY_ remains much less than 1 until the saturation of the receptor has occurred. The rationale behind setting (1/Kd_YZ_)·(β_2_/α_2_) equal to 10^4^ is given in the Methods (part 1a).

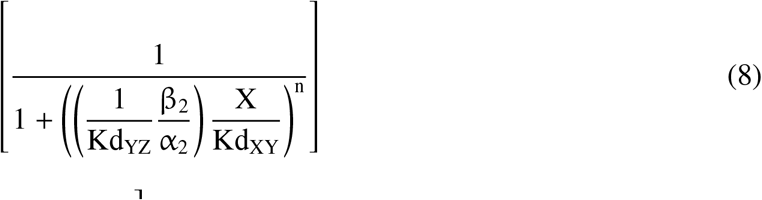

**Figure 3:**
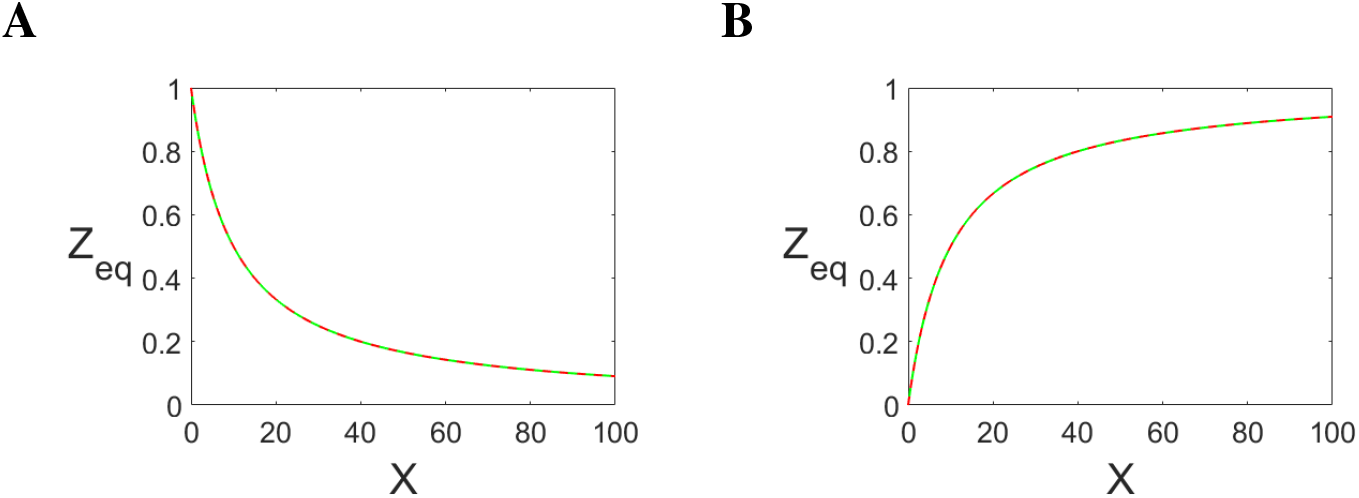
(**A**) The green curve represents the expression 1/(1 + 10^4^((X/10^5^)/(1 + (X/10^5^)))). The red curve: 1/(1 +(X/10)). (**B**) The green curve represents the expression 10^4^((X/10^5^)/(1 + (X/10^5^)))/(1 + 10^4^((X/10^5^)/(1 + (X/10^5^)))). The red curve: (X/10)/(1 + (X/10)).

**Figure 4:**
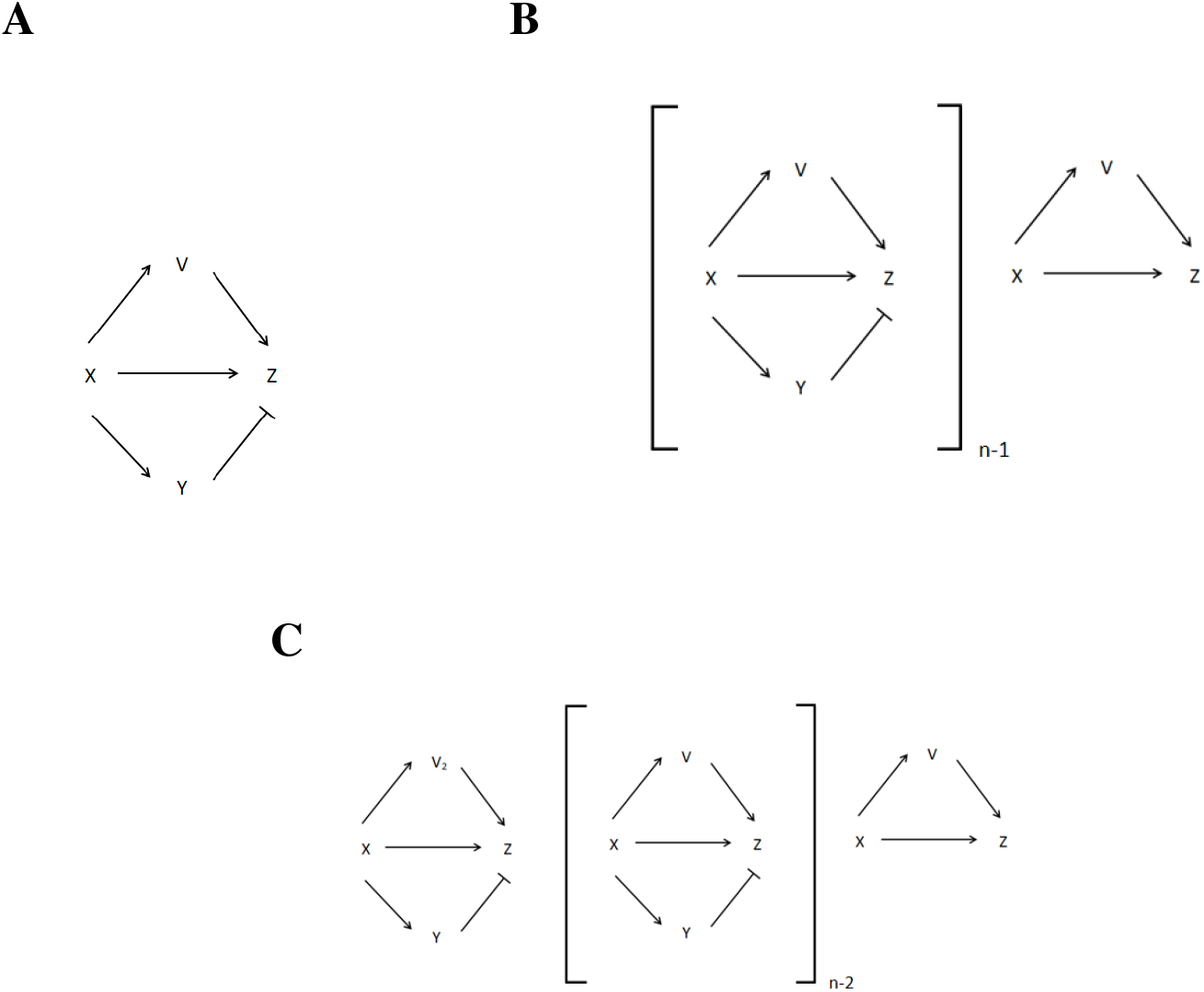
(**A**) FNM whose series can generate linear input-output relationship and compute logarithm. (**B**) The series for generating linear input-output relationship. (**C**) The series for computing logarithm.

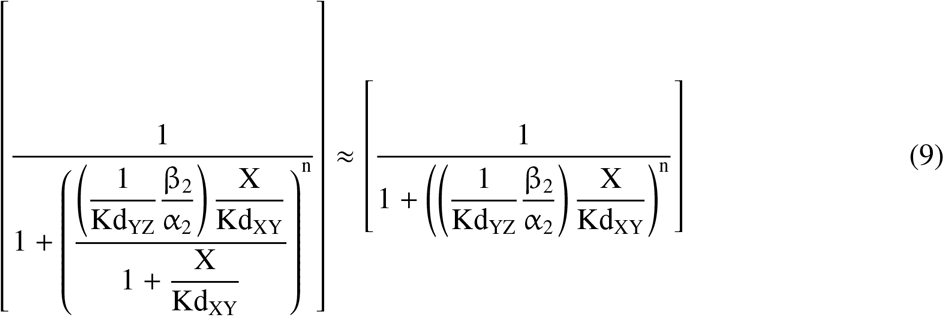

After applying the approximation, eq. 7 becomes

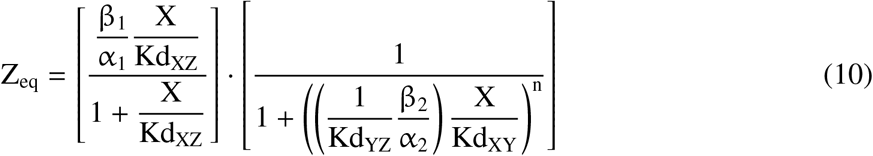

To fulfill the aforementioned demand on the value of parameters of the negative arm, Kd_YZ_ · (α_2_/β_2_) · Kd_XY_, the effective equilibrium dissociation constant of this arm, is set equal to Kd_XZ_. The total output from the five receptors is

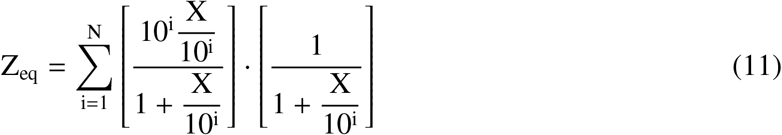

The following is non-approximated form of the above eq.

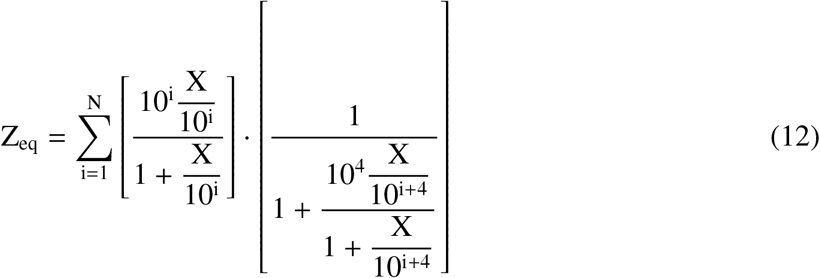

However, an input signal can generate output Z from those receptors also whose Kd_XZ_ is larger than the input signal by more than one order of magnitude. In the present case, this means that the input signal of 10^2^ arb. units being able to generate the output from the receptors whose Kd_XZ_ is equal to 10^4^ and 10^5^ arb. units, respectively. Output Z_eq_ generated by each of the two receptors is approximately equal to 10^2^ arb. units, which is of the same order as Z_eq_ generated by the receptor whose Kd_XZ_ is equal to 10^3^ arb. units. The output from the receptors whose Kd_XZ_ is larger than the input signal by more than one order of magnitude is removed by adding to the motif a “positive arm”: the input signal X engenders production of the molecule V, which must also be bound along with X to the producer element of Z in order to engender Z production. Parameters of the positive arm are such that it lets mainly those receptors generate the output whose Kd_XZ_ is at most one order of magnitude larger than the input signal.

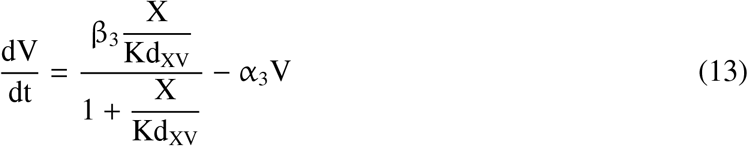

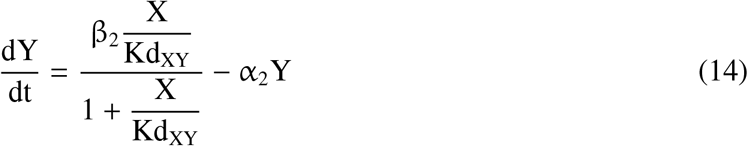

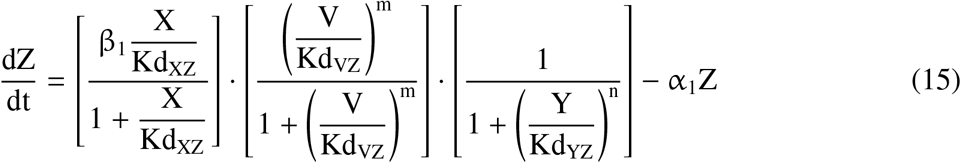

where, β_1_, β_2_, and β_3_ are the rate constants for production of Z, Y, and V, respectively, by X. α_1_, α_2_, and α_3_ are the degradation rates of Z, Y, and V, respectively. Kd_XZ_, Kd_XY_, and Kd_XV_ are the equilibrium dissociation constants for the binding of X to the element of Z, Y, and V, respectively, which causes production of Z, Y, and V, respectively. Kd_YZ_ and Kd_VZ_ are the equilibrium dissociation constants for the binding of Y and V, respectively, to the element of Z, which causes Z production.

The equilibrium value of V, Y, and Z

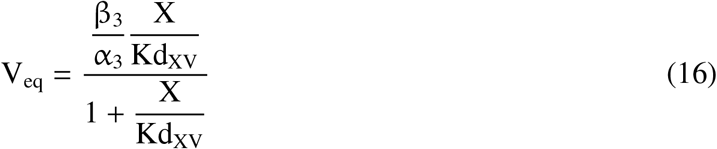

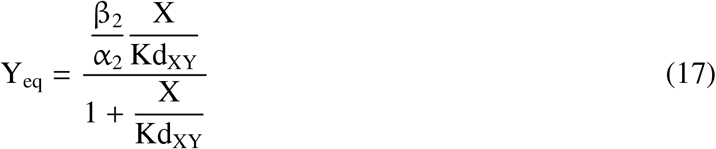

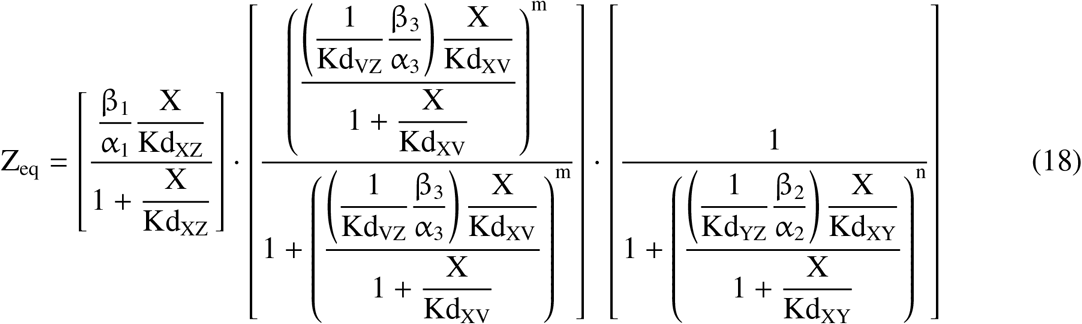

The formulation demands that the expression in the second pair of brackets on the RHS of eq. 18, representing the generation of output Z_eq_ by the input signal X and the molecule V according to the AND logic, should be of the form X^n^/(1 + X^n^). This is achieved in the following way. The expression in the second pair of brackets on the RHS of eq. 18 is approximately equal to the expression 19 (which is of the required form), at least in the parameter range of interest (i.e., until X is not at least one order of magnitude larger than Kd_XZ_), if Kd_XV_ ≫ Kd_XZ_ and the order of magnitude of (1/Kd_VZ_) · (β_3_/α_3_) is at least four (Figure 3B). This is because when Kd_XV_ ≫ Kd_XZ_, X/Kd_XV_ remains much less than 1 until the saturation of the receptor has occurred. Again, the reasoning behind setting (1/Kd_VZ_) · (β_3_/α_3_)) equal to 10^4^ is given in the Methods (part 1a).

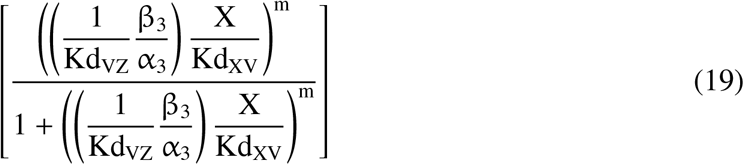

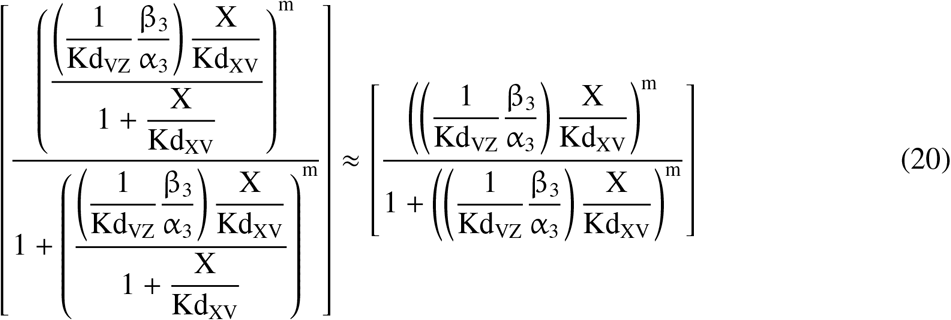

After applying the approximation, eq. 18 becomes

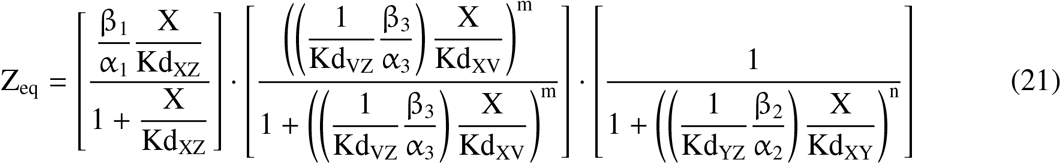

To fulfill the aforementioned demand on the value of parameters of the positive arm, Kd_VZ_ · (α_3_/β_3_)· Kd_XV_, the effective equilibrium dissociation constant of this arm, is set equal to Kd_XZ_/10. The total output from the five receptors is

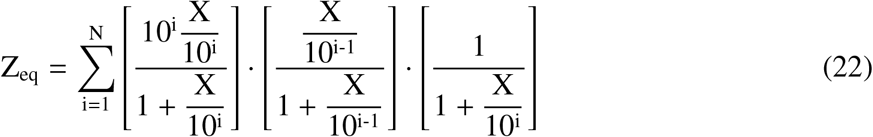

The following is non-approximated form of the above eq.

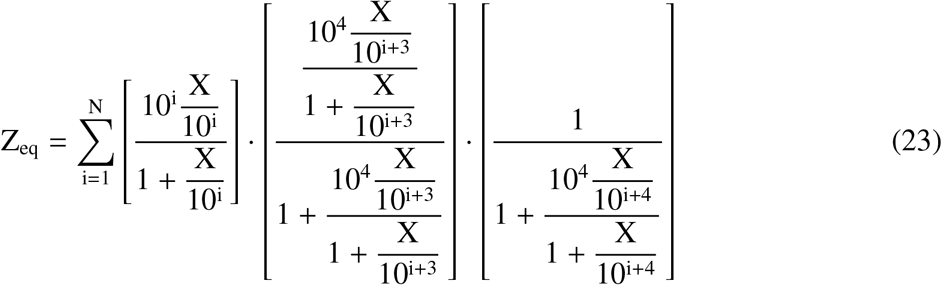

Hence, together the negative arm and the positive arm create a window which show maximum response for an input signal of a particular order of magnitude and whose output declines progressively as the distance between the preferred order of magnitude and the order of magnitude of input signal increases in either direction. Further, the input-output curve has positive skewness. Figure 5 shows input-output relationship of a single four-node motif (FNM) with four combinations of the Hill-coefficients. Increasing the value of m introduces steepness at the leftmost end of the output, whereas increasing the value of n causes the output to have higher and sharper peak and decline quickly.

**Figure 5:**
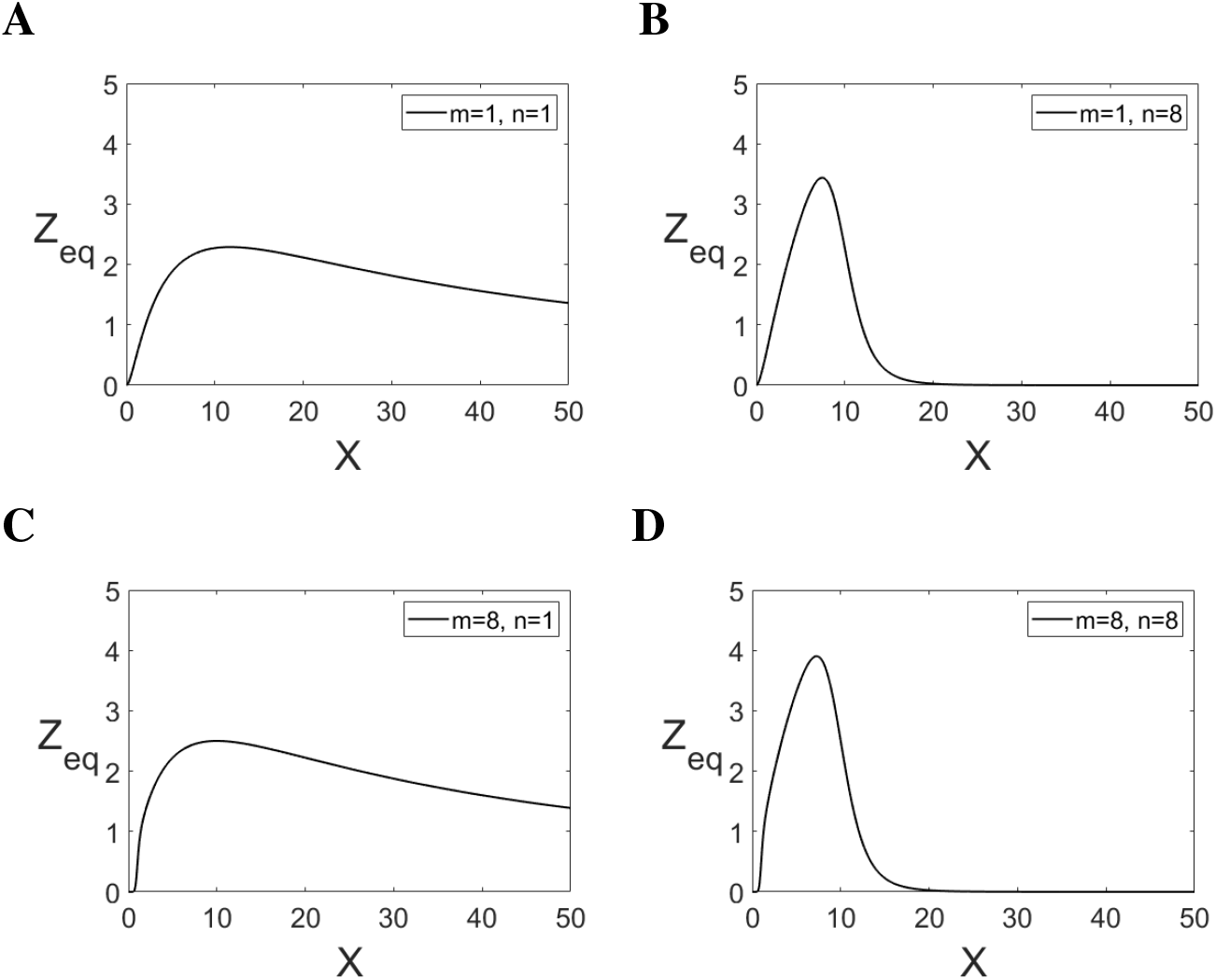
**(A-D)** The output of FNM defined by the eq. [10 · (X/10)/(1 + X/10)] · [(X/1)^m^/(1 + (X/1)^m^)] · [1/(1 + (X/10)^n^)] with different values of the Hill-coefficients m and n.

Notably, because a) β_1_/α_1_ is equal to Kd_XZ_, b) Kd_XY_=10^4^·Kd_XZ_, c) Kd_XV_=10^3^·Kd_XZ_, d) (1/Kd_YZ_)· (β_2_/α_2_) = (1/Kd_VZ_)·(β_3_/α_3_) = 10^4^, the value of Kd_XZ_ is sufficient to specify all other parameters of FNM whose series produces linear input-output relationship. The last receptor should not have the negative arm, otherwise a very large input signal will generate a very small output. Thus, the eq. for the output should be:-

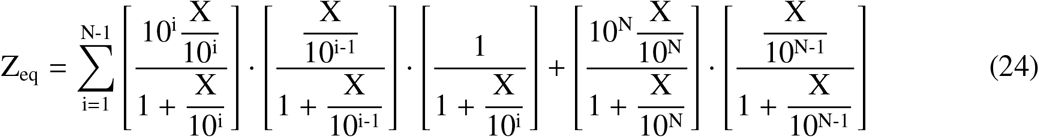

The following is non-approximated form of the above eq.

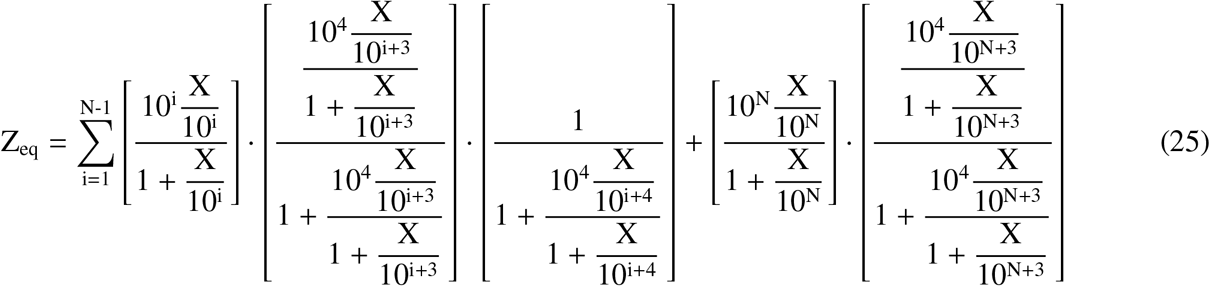

How well the expression on the RHS of eq. 21 approximates the expression on the RHS of eq. 18 was determined by analyzing how well the first term on the RHS of eq. 22 approximates the first term on the RHS of eq. 23, using linear regression (see Methods, part 2a). As stated above, in order to provide sensitive detection, ideal relationship between an input signal and the output should be a straight line with the slope equal to 1 and the y-intercept equal to 0. Thus, in order to assess if, given a range of Kd_XZ_, there is a range of input signal in which the series of FNM could generate linear input-output relationship, straight line was fitted with output Z_eq_ given by eq. 25 (the non-approximated form) for various ranges of Kd_XZ_ and input signal (Table 1). See Methods (part 3) for details of the fitting procedure.

**Table 1:** Slope and goodness of fit (R^2^) of linear regression of output Z_eq_ against the input values for various ranges of signal and Kd_XZ_.

| Signal Range (across)<br>$Kd_{XZ}$ Range (down) | $10^0-10^2$ | $10^{-1}-10^3$ | $10^{-2}-10^4$ | $10^{-3}-10^5$ | $10^{-4}-10^6$ |
| --- | --- | --- | --- | --- | --- |
| $10^{-2}-10^4$ | 0.7383<br>(1.0000) | 0.6891<br>(1.0000) | 0.4606<br>(0.9976) | NA | NA |
| $10^{-3}-10^5$ | 0.7477<br>(1.0000) | 0.7386<br>(1.0000) | 0.6896<br>(1.0000) | 0.4618<br>(0.9976) | NA |
| $10^{-4}-10^6$ | 0.7487<br>(1.0000) | 0.7477<br>(1.0000) | 0.7387<br>(1.0000) | 0.6899<br>(1.0000) | 0.4625<br>(0.9976) |
| $10^{-5}-10^7$ | 0.7489<br>(1.0000) | 0.7488<br>(1.0000) | 0.7478<br>(1.0000) | 0.7387<br>(1.0000) | 0.69<br>(1.0000) |
| $10^{-6}-10^8$ | 0.7489<br>(1.0000) | 0.7489<br>(1.0000) | 0.7488<br>(1.0000) | 0.7478<br>(1.0000) | 0.7388<br>(1.0000) |

For the middle intervals, the slope of the input-output relationship converges to the value 0.7489 when the input signal is a multiple of 10. However, the limiting values of the slope are not identical when the input signal is not powers of 10 (Figure 8A). The percentage difference between the maximum value and the minimum value of the slope is 0.8250% and 0.8318%, depending on whether the difference is normalized by the maximum value or the minimum value of the slope, respectively. Although the slope of the line given by eq. 24/25 oscillates between 0.7489 (when X is a multiple of 10) and 0.7551 (when X is 3 times the multiple of 10), the difference is so small that the input-output relationship can be considered to be quasi-linear, having the average slope of 0.75136. Further, the same results as presented in table 1 were got when the range of Kd_XZ_ and concomitantly the range of input signal was shifted by the factor of 0.01, 0.1, 10, and 100 (see Methods, part 3). However, if the number of receptors in a series were such that, for a given range of input signal, the slope had reached its limiting value, then as the range of Kd_XZ_ and input signal were multiplied by the factor of 0.01, 0.1, 10, and 100, the y-intercept of the fitted line turned out to be 0.01, 0.1, 10, and 100 times the previous y-intercept, respectively. Thus, although the slope remains constant, the y-intercept increases to 10^k^ (where k is a scalar) its previous value if the multiplicative factor, both for the range of Kd_XZ_ and input signal, is 10^k^ with k>1. However, if the modified series (the series whose range of Kd_XZ_ is multiplied by 10^k^ with k>1) is extended at both ends by k receptors, the y-intercept goes back to its previous value. In general, if the number of receptors in a series are such that, for a given range of input signal, the slope has reached its limiting value, then for the range of input signal, the y-intercept reduces to 10^-k^ its value if the series is extended in both directions by k receptors. Thus, the y-intercept can be reduced to arbitrary low values.

The average y-intercept for the ten values of X is 2.1601e-04 with the standard deviation equal to 3.3083e-08. Taken together, if output Z_eq_ of this series is multiplied by the reciprocal of 0.75136, one gets an almost straight line passing through the origin and having the slope 1. Because there is no limit on the number of the receptors, the series of FNM can generate quasi-linear input-output relationship over arbitrary range of input signal. Note that in order to provide the quasi-linear input-output relationship over the range 10^a^-10^b^ (where, a < b and both are integers), the series of receptors whose Kd_XZ_ ranges at least from 10^a-5^ to 10^b+5^ is required.

For the values of m and/or n being more than one, the output becomes oscillatory around the output generated with m=1 and n=1 (Figure 6). This implies that V and Y molecules should be present as monomers in a real system. Arguably, the output produced by eq. 24/25 needs to be decoded by the brain in order to infer the input signal.

**Figure 6:**
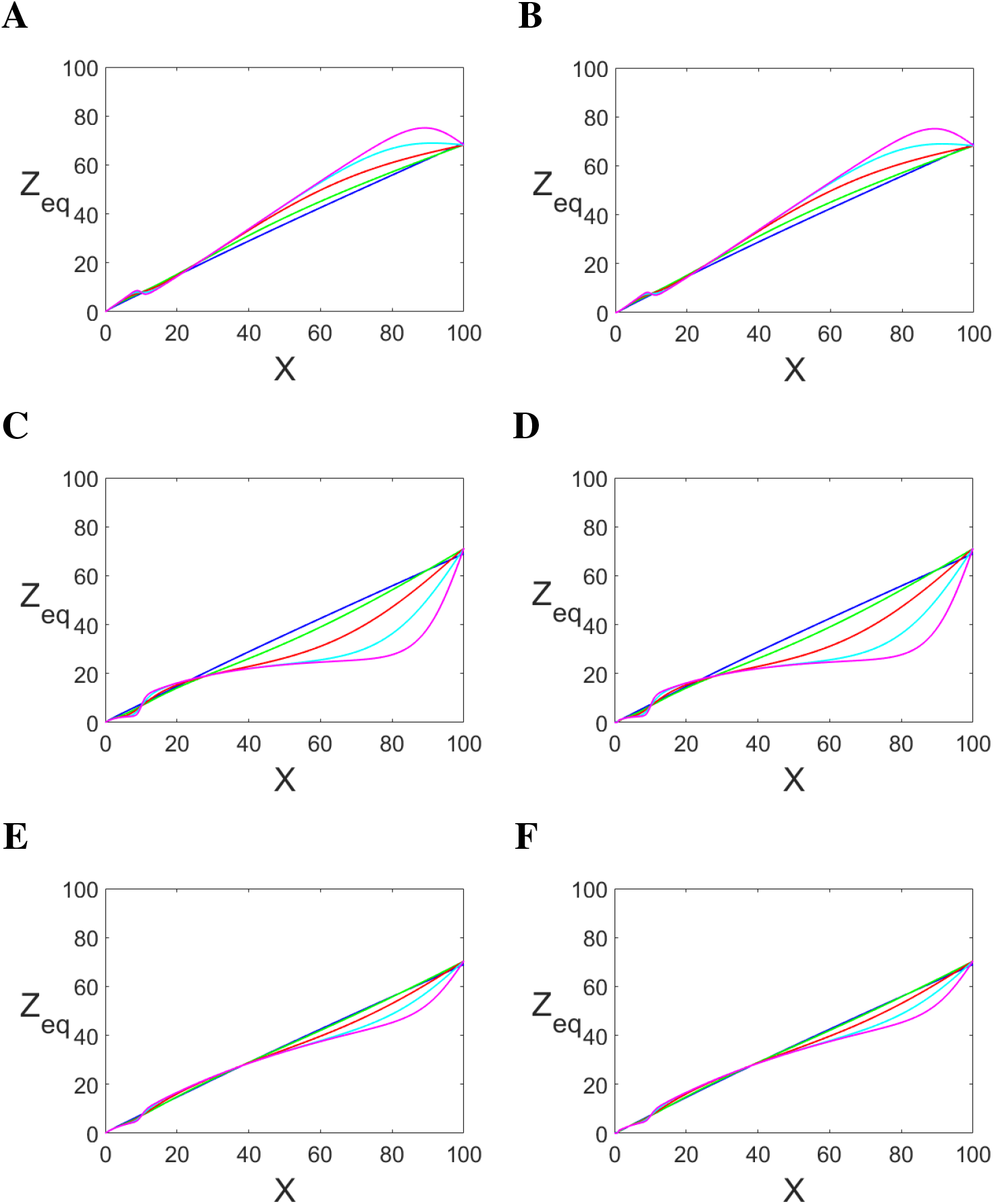
The total output (Z_eq_) from the first three FNM receptors for the approximated form (**A**) and non-approximated form (**B**) when m=1, n=1 (blue); m=1, n=2 (green); m=1, n=4 (red); m=1, n=8 (cyan); m=1, n=16 (pink). The total output (Z_eq_) from the first three FNM receptors for the approximated form (**C**) and non-approximated form (**D**) when m=1, n=1 (blue); m=2, n=1 (green); m=4, n=1 (red); m=8, n=1 (cyan); m=16, n=1 (pink). The total output (Z_eq_) from the first three FNM receptors for the approximated form (**E**) and non-approximated form (**F**) when m=1, n=1 (blue); m=2, n=2 (green); m=4, n=4 (red); m=8, n=8 (cyan); m=16, n=16 (pink).

**Figure 7:**
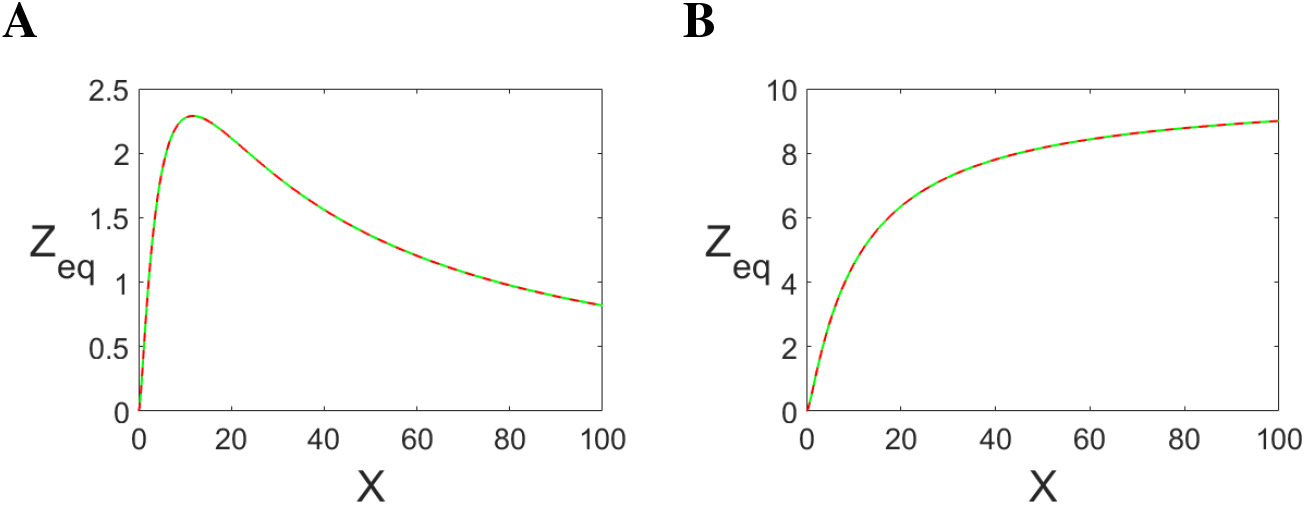
(**A**) The green curve and red curve represent the first term on the RHS of eq. 23 and eq. 22, respectively. (**B**) The green curve and red curve represent the first term on the RHS of eq. 29 and eq. 28, respectively.

**Figure 8:**
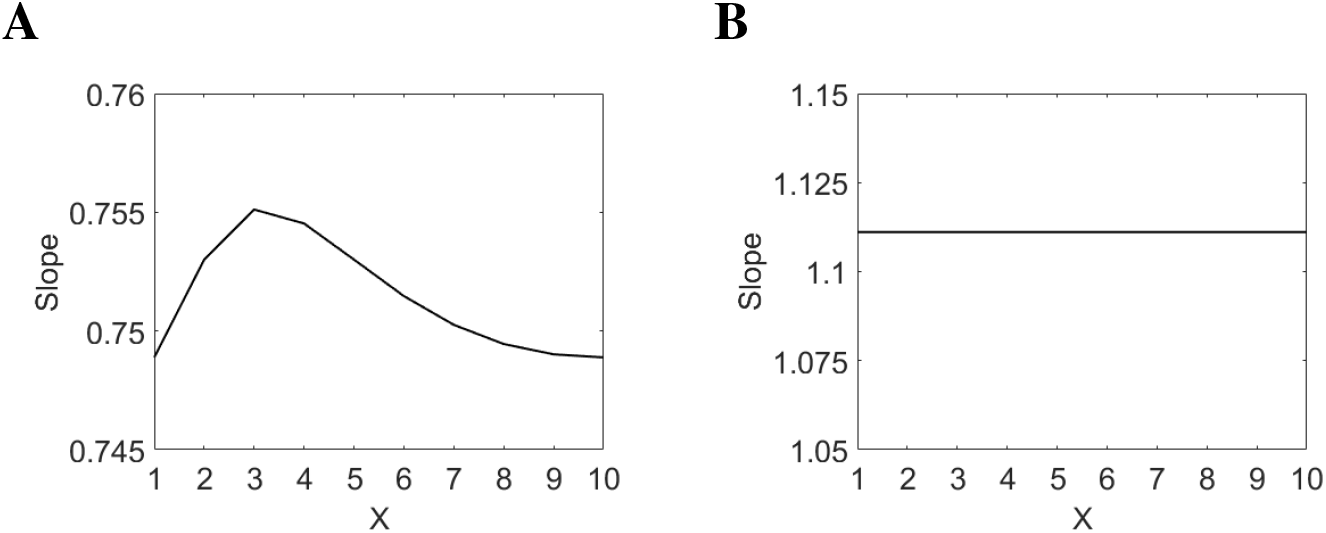
The limiting value of the slope from the linear regression of Z_eq_ against the input signal’s value (X·10^k^) for the series of FNM (**A**) and the series of C1-FFL (**B**). Note that the limiting value of the slope for the two series is equal to 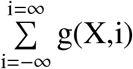 and 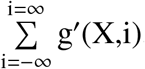, respectively

**Figure 9:**
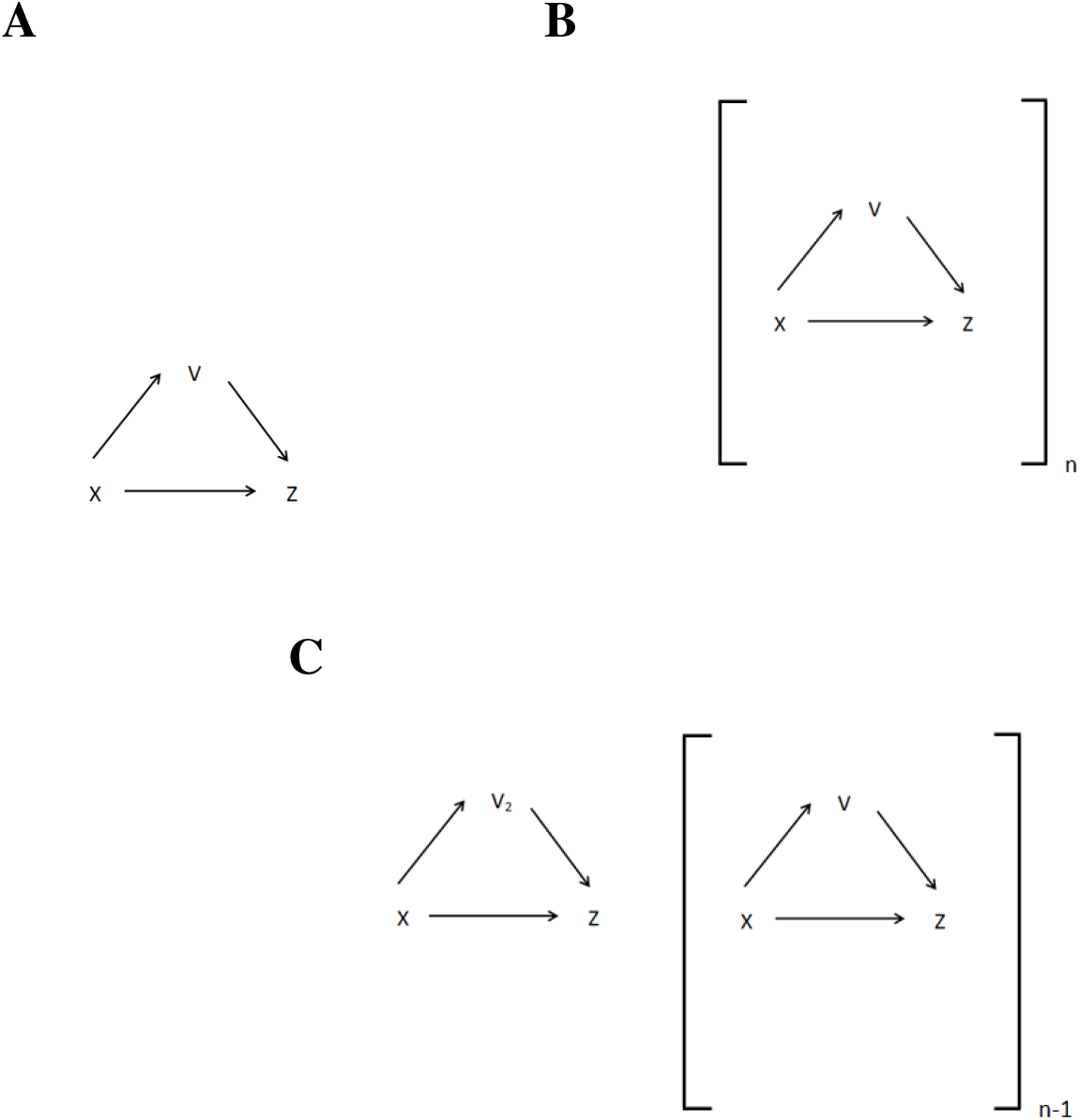
(**A**) The C1-FFL motif whose series can generate linear input-output relationship and compute logarithm. (**B**) The series for generating linear input-output relationship. (**C**) The series for computing logarithm.

**Figure 10:**
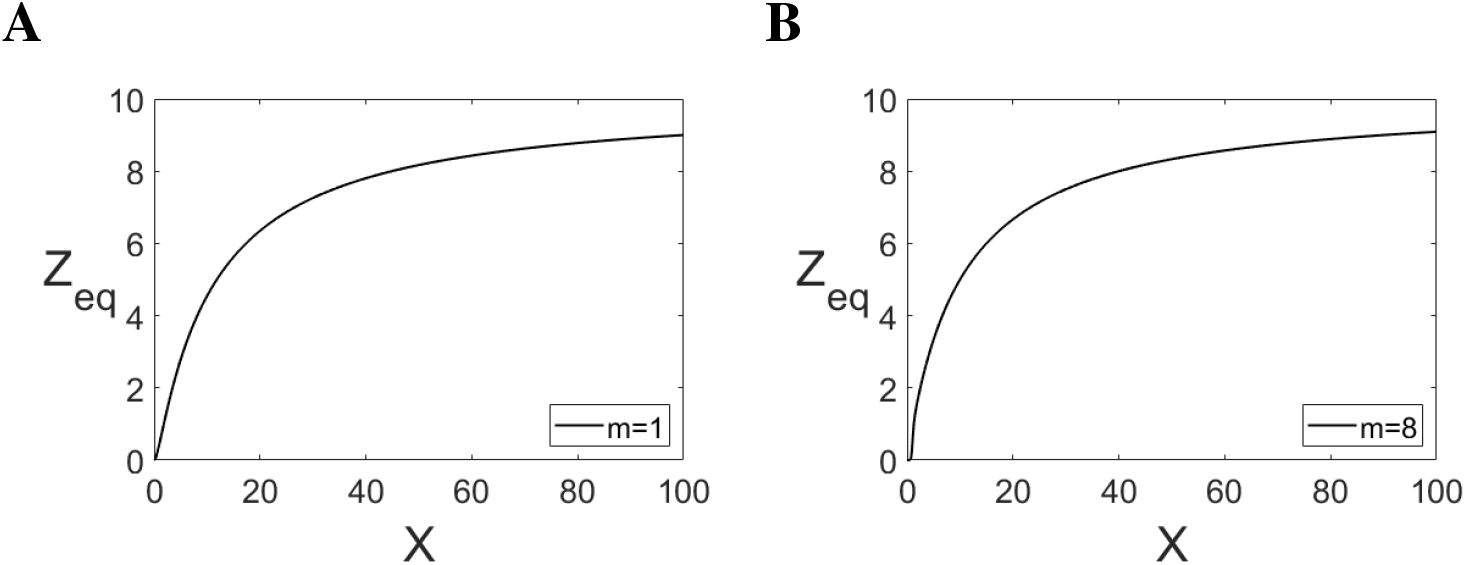
(**A-D**) The output of FNM defined by the eq. [10·(X/10)/(1 + X/10)]·[(X/1)^m^/(1 + (X/1)^m^)] with different combinations of the value of the Hill-coefficients m and n.

### 2.2. The Mathematical Basis of the Quasi-Linear Input-Output Relationship Provided by the Series of the Four-node Motif

Equation 24 provides quasi-linear input-output relationship. It differs from eq. 22 in that its last receptor lacks the negative arm. However, this difference is inconsequential if a large number of receptors flank the receptor which contributes the most to output Z_eq_ (i.e., whose Kd_XZ_ is an order of magnitude higher than the input signal’s value), as then output Z_eq_ of the last receptor is negligible. In the analysis that follows, because the series of infinite FNM receptors is considered, the output of eq. 22 and eq. 24 are practically identical, and thus eq. 22 is considered. Equation 22 is equivalent to Z_eq_ = m·X (because the y-intercept is negligible) and can be written as

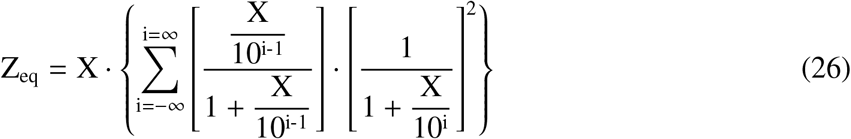

Equation 26 can be abbreviated in the following way:

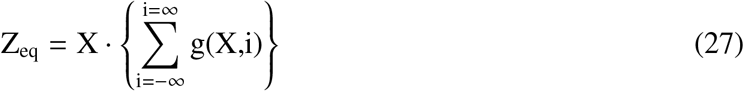

Now it remains to be shown that the series in the curly brackets of eq. 26/27 converges in both negative and positive direction to a value which, given an input value X, remains the same for 10^k^·X (where is k is an integer). For very large positive values of i, g(X,i) ≈ X/10^i-1^, and for very large negative values of i, g(X,i) ≈ (10^i^/X)^2^. The two asymptotic forms of g(X,i) are convergent series, as the first one is the geometric series and the second one is the geometric series having only the even powers. Now it is shown that given an input signal X, Z_eq_ remains the same for 10^k^·X (k being an integer). Replacing X by 10^k^·X and doing the transformation i → i+k leaves g(X,i) in eq. 26/27 unchanged. However, because i runs from minus infinity to plus infinity, the transformation of i → i+k does not make any difference. Substituting X/10^i^ with Y in eq. 26, and writing g(X,i) as p(Y)/q(Y), it can be seen that 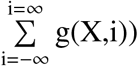 converges in both directions when the degree of q(Y) is at least one more than the degree of p(Y). This means that g(X,i) of the form [(X/10^i-1^)/(1+X/10^i-1^)]·[1/(1+X/10^i^)] (hereinafter called g (X,i)) should also be able to provide at least a quasi-linear input-output relationship. This is indeed the case: for very large positive values of i, g (X,i) ≈ X/10^i-1^, and for very large negative values of i, g (X,i) ≈ (10^i^/X). Both are geometric series and hence convergent. Note that g (X,i) corresponds to coherent type-1 feed-forward loop (C1-FFL); that is, FNM minus the negative arm. The following is the eq. for the output of the series of C1-FFL receptors.

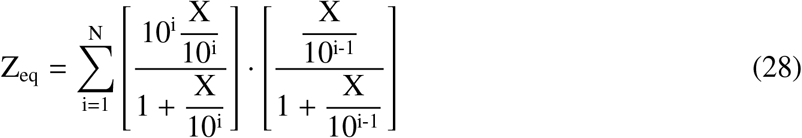

The following is non-approximated form of the above eq.

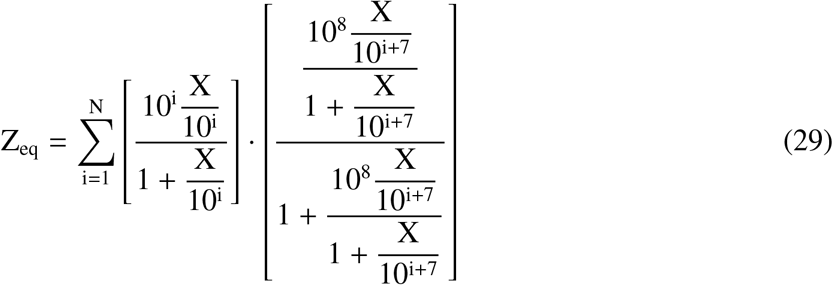

Note that (1/Kd_YZ_)·(β_2_/α_2_) and (1/Kd_VZ_) · (β_3_/α_3_)) are set equal to 10^8^ (unlike in eq. 23/25, where the two quantities are set equal to 10^4^). The rationale behind setting the two quantities equal to 10^8^ is given in the Methods (part 1b). As with FNM given in eq. 23/25, the value of Kd_XZ_ is sufficient to specify all other parameters of C1-FFL whose series generates linear input-output relationship. This is because a) β_1_/α_1_ is equal to Kd_XZ_, b) Kd_XV_=10^7^·Kd_XZ_, c) (1/Kd_YZ_) · (β_2_/α_2_) = (1/Kd_VZ_) · (β_3_/α_3_) = 10^8^.

Like for FNM, how well the first term on the RHS of eq. 28 approximates the first term on the RHS of eq. 29 was determined using linear regression (see Methods, part 2b). As stated above, in order to provide sensitive detection, ideal relationship between an input signal and the output should be a straight line with the slope equal to 1 and the y-intercept equal to 0. Thus, as previously, in order to assess if, given a range of Kd_XZ_, there is a range of input signal in which the series of C1-FFL could generate linear input-output relationship, straight line was fitted with output Z_eq_ given by eq. 29 (the non-approximated form) for various ranges of Kd_XZ_ and input signal (Table 2). See Methods (part 3) for details of the fitting procedure.

**Table 2:** Slope and goodness of fit (R^2^) of linear regression of output Z_eq_ against the input values for various ranges of signal and Kd_XZ_.

| **Signal Range (across)**<br>**$Kd_{XZ}$ Range (down)** | $10^0-10^2$ | $10^{-1}-10^3$ | $10^{-2}-10^4$ | $10^{-3}-10^5$ | $10^{-4}-10^6$ |
| --- | --- | --- | --- | --- | --- |
| $10^{-2}-10^4$ | 1.1<br>(1.0000) | 1.008<br>(0.9999) | 0.5513<br>(0.9933) | NA | NA |
| $10^{-3}-10^5$ | 1.11<br>(1.0000) | 1.1<br>(1.0000) | 1.009<br>(0.9999) | 0.5535<br>(0.9933) | NA |
| $10^{-4}-10^6$ | 1.111<br>(1.0000) | 1.11<br>(1.0000) | 1.1<br>(1.0000) | 1.01<br>(0.9999) | 0.5549<br>(0.9933) |
| $10^{-5}-10^7$ | 1.111<br>(1.0000) | 1.111<br>(1.0000) | 1.11<br>(1.0000) | 1.1<br>(1.0000) | 1.01<br>(0.9999) |
| $10^{-6}-10^8$ | 1.111<br>(1.0000) | 1.111<br>(1.0000) | 1.111<br>(1.0000) | 1.11<br>(1.0000) | 1.1<br>(1.0000) |

For the middle intervals, the slope of input-output relationship generated by the series of C1-FFL converges to the value 1.111 (or 10/9). Also, the slope is independent of the input signal’s value (Figure 8B). Further, the same results as presented in table 2 were got when the range of Kd_XZ_ and concomitantly the range of input signal were shifted by the factor of 0.01, 0.1, 10, and 100 (see Methods, part 3). However, if the number of receptors in a series were such that, for a given range of input signal, the slope had reached its limiting value, then as the range of Kd_XZ_ and input signal were multiplied by the factor of 0.01, 0.1, 10, and 100, the y-intercept of the fitted line turned out to be 0.01, 0.1, 10, and 100 times the previous y-intercept, respectively. Thus, although the slope remains constant, the y-intercept increases to 10^k^ (where k is a scalar) its previous value if the multiplicative factor, both for the range of Kd_XZ_ and input signal, is 10^k^ with k>1. However, if the modified series (the series whose range of Kd_XZ_ is multiplied by 10^k^ with k>1) is extended at both ends by k receptors, the y-intercept goes back to its previous value. In general, if the number of receptors in a series are such that, for a given range of input signal, the slope has reached its limiting value, then for the range of input signal, the y-intercept reduces to 10^-k^ its value if the series is extended in both directions by k receptors. Thus, the y-intercept can be reduced to arbitrary low values.

The average y-intercept for the ten values of X is 2.1592e-04 with the standard deviation equal to 3.3091e-08. Taken together, if output Z_eq_ of this series is multiplied by the reciprocal of 1.111, one gets a straight line passing through the origin and having the slope 1. Because there is no limit on the number of the receptors, the series of C1-FFL can generate perfectly linear input-output relationship over arbitrary range of input signal. Note that in order to provide the linear input-output relationship over the range 10^a^-10^b^ (where, a < b and both are integers), the series of receptors whose Kd_XZ_ ranges at least from 10^a-4^ to 10^b+4^ is required. For the values of m and/or n being more than one, the output becomes oscillatory around the output generated with m=1 and n=1 (Figure 11). This implies that V and Y molecules should be present as monomers in a real system. As with the series of FNM, the output produced by eq. 28/29 needs to be decoded by the brain in order to infer the input signal.

**Figure 11:**
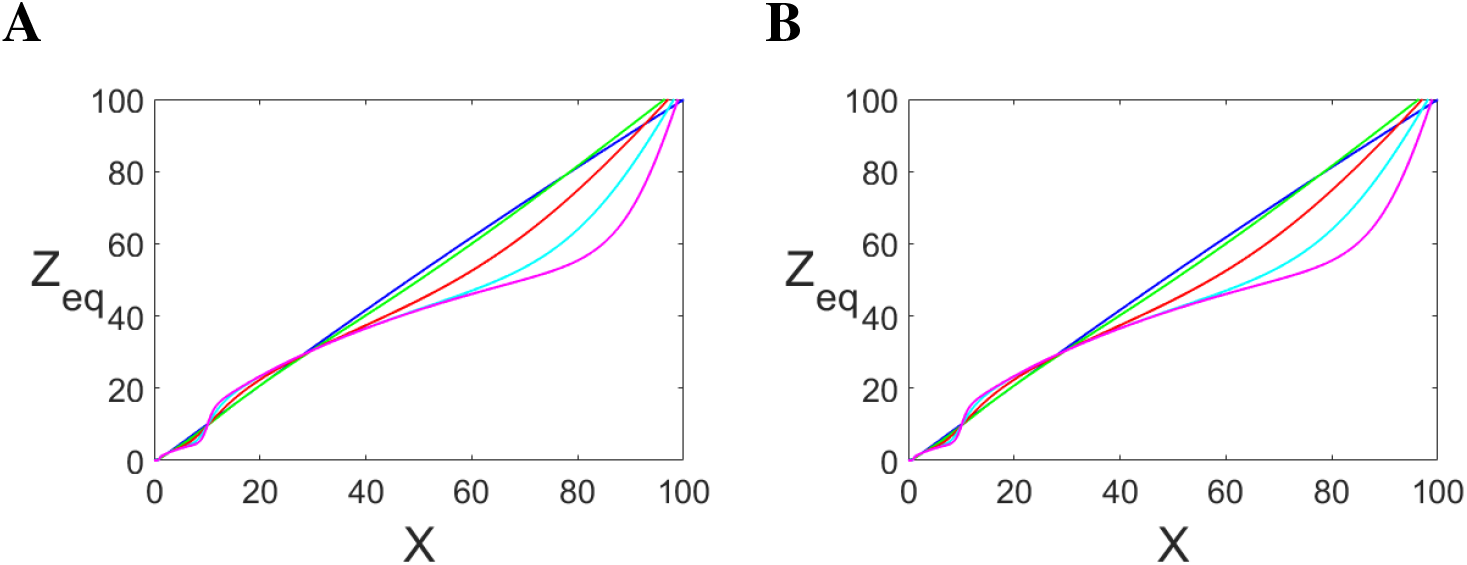
The total output (Z_eq_) from the first three C1-FFL receptors for the approximated form (**A**) and non-approximated form (**B**) when m=1 (blue); m=2 (green); m=4 (red); m=8 (cyan); m=16 (pink).

### 2.3. Weber’s Law

Now I present a formalism to get Weber’s law from the series of FNM and the series of C1-FFL. Equation 24/25, when multiplied by the factor of 1/0.75136, yields an input-output relationship which, to a very good approximation, is linear for intermediate range of input signal (X); that is, Z_eq_ ≈ X. However, as shown above, because the deviation from linearity is minuscule, the input-output relationship is assumed to be linear (i.e., Z_eq_ = X) for the purpose of the analysis below. On the other hand, eq. 28/29, when multiplied by the factor of 1/1.111 (or 10/9), yields a perfectly linear input-output relationship for intermediate range of X; that is, Z_eq_ = X. The following analysis is common for both the series. Given an input signal (X = µ), the output (Z_eq_) is considered to be the result of sampling from a Gaussian distribution with mean equal to µ and standard deviation linearly proportional to µ; that is, σ = fµ, where f is the Weber fraction, the ratio of the smallest perceptual change in an input signal (ΔX_min_) and the background signal (X_background_): ΔX_min_/X_background_ = f. It is posited here that the subject is able to perceive a change in the input signal when the ratio of the non-overlapping region, towards the right, between the two Gaussians (shaded region in figure 12) and the area under the Gaussian centered on µ_1_ is above a certain threshold value. Because the Gaussians in the consideration are normalized, the ratio is equal to the non-overlapping region on the right, R.

**Figure 12:**
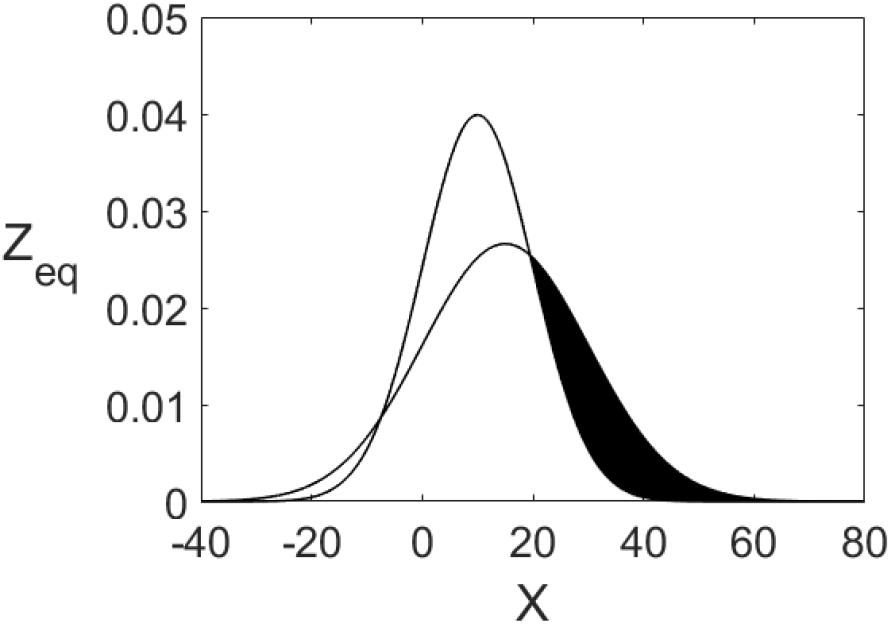
The Gaussian on the left has mean µ_1_=10, and σ_1_=10, and the Gaussian on the right has mean µ_2_=15 and σ_2_=15. The shaded portion is the non-overlapping region on the right, R

As is defined now, the region R is just below the threshold value when µ_2_ − µ_1_ = σ_1_; that is, when the difference between the mean of the current input signal and the mean of the background signal is equal to the standard deviation of the background signal. In other words, σ_1_ is the just noticeable difference (JND). The region R is F_1_(p_2_) - F_2_(p_2_), where F(X) is the cumulative distribution function of Gaussian distribution and p_2_ is the farther point of intersection of the two Gaussians when µ_2_ −µ_1_ = σ_1_. That is, p_2_ is the bigger-valued solution of the quadratic eq. below. Note that the total output Z_eq_, given by eq. 24/25, is the sum of Z_eq_ of individual receptors. But that still does not affect the formalism above because the individual outputs are independent, and thus the sum of the individual means is µ and the sum of the individual standard deviations is σ. This is further corroborated by the stochastic simulation later below

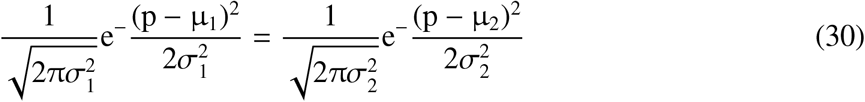

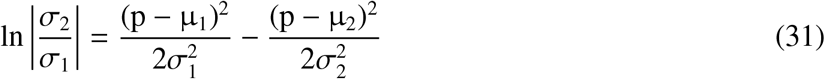

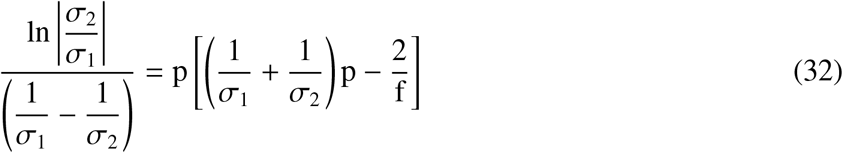

Setting 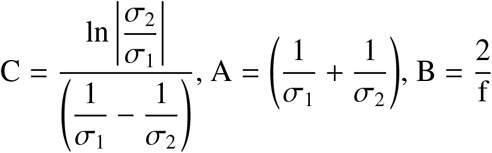 eq. 32 becomes

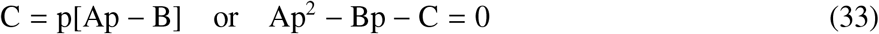

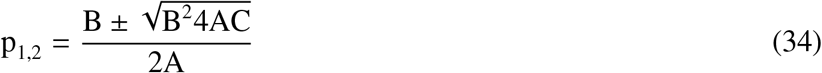

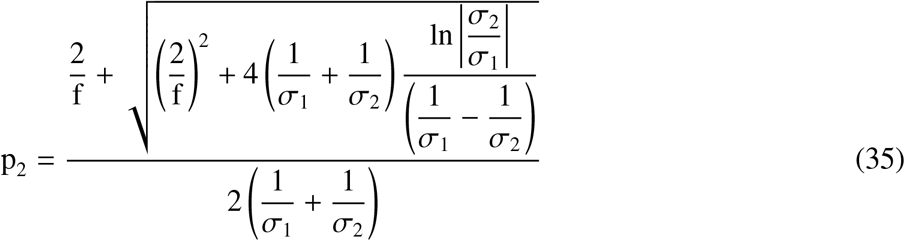

Setting 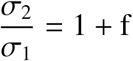

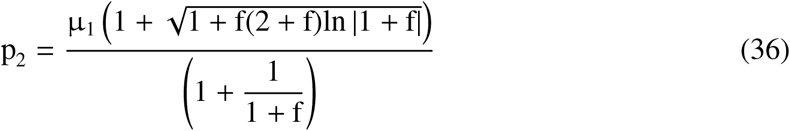

The eq. 36 can be written as p_2_ = µ_1_g(f)

The CDF of a Gaussian distribution with mean µ and variance σ^2^ is

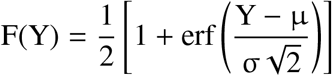

Hence

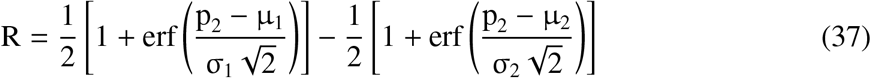

Inserting the value of p_2_ in eq. 37

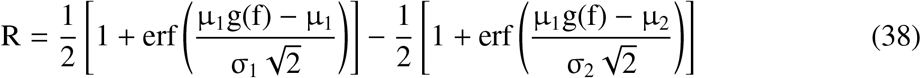

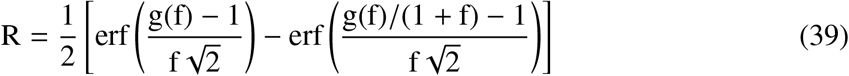

Being a function of f alone, R can be written as R = h(f). It also implies that the value of R remains the same when the two means are multiplied by a scalar. This is because the two Gaussians become flatter and the point p_2_ goes farther. Hence, the region R stays constant when the two signals are multiplied by a scalar, which is Weber’ law. In the range of f from 0 to 1, R is a monotonically decreasing function of f (data not shown); hence h^−1^R is also a function.

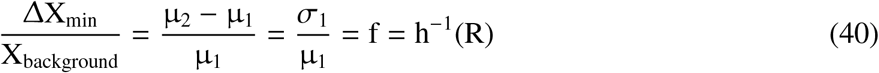

In the formalism presented here (i.e., the linear model), Weber’s law is the result of a Gaussian internal noise whose standard deviation is proportional to the input signal. On the other hand, Fechner’s explanation is based on the assumption that the output is proportional to a logarithm of the input signal, and JND therefore is a function of a constant increment in the output (eqs. 44 and 45).

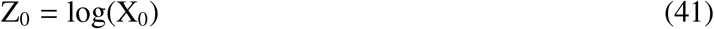

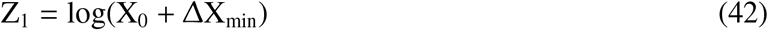

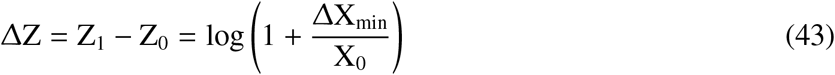

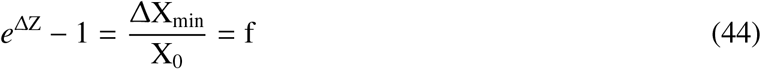

For ΔZ ≪ 1, eq. 44 becomes

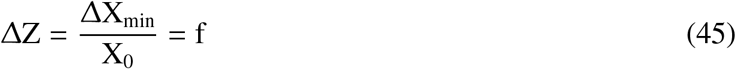

However, Fechner’s formulation does not provide a biological basis of the relation between the Weber fraction and ΔZ (eqs. 44 and 45). Also, the output being logarithm of the input signal does not seem natural. Two Wildebeests butting their head against each other should not compute logarithm of the force that they receive from the other in the process. Otherwise, a force four times as large will appear larger by only log4. With such a sensory system the weaker animal will be much more injured at the end of the fight than if the sensory system converted the input signal to an equally valued output.

The formalism for Weber’s law presented in this paper assumes that output Z_eq_ given by eq. 25 (approximated as eq. 24) and eq. 29 (approximated as eq. 28), respectively, to be noisy and thus considers the output to be a random sample from a Gaussian distribution whose mean is the input signal’s value and the standard deviation is the Weber fraction times the mean. In order to validate this assumption, output Z_eq_ of eq. 25 and eq. 29, respectively, should be shown to be a Gaussian distribution with mean equal to a constant times the input signal’s value and standard deviation equal to a constant times the Weber fraction times the mean when the output generated by individual components of FNM and C1-FFL, respectively, is a random sample from their corresponding Gaussian distributions (See Methods, part 4).

For the series of FNM: for the values of f up to 0.2, the average slope of the fitted line –for the mean– was consistently 1.45 (standard deviation being of the order of magnitude -5 at worst), the average y-intercept of the fitted line –for the mean– was very close to zero; that is, being of the order of magnitude -4 at worst (with standard deviation being of the order of magnitude -5 at worst). The average slope of the fitted line –for the standard deviation– was consistently 1.39 (standard deviation being of the order of magnitude -3 at worst), the average y-intercept of the fitted line –for the standard deviation– was very close to zero; that is, being of the order of magnitude -4 at worst (with standard deviation being of the order of magnitude -5 at worst). And the average R^2^ was equal to 1 (that is, standard deviation being zero) for both the mean and standard deviation. For higher values of f (i.e., 0.5 and 1), the fit became progressively poor (Supplementary Information, Table 8).

For the series of C1-FFL: for the values of f up to 0.2, the average slope of the fitted line –for the mean– was consistently very close of 0.9 (standard deviation being of the order of magnitude -4 at worst), the average y-intercept of the fitted line –for the mean– was very close to zero; that is, being of the order of magnitude -3 at worst (with standard deviation being of the order of magnitude -5 at worst). The average slope of the fitted line –for the standard deviation– was consistently very close of 1.26 (standard deviation being of the order of magnitude -4 at worst), the average y-intercept of the fitted line –for the standard deviation– was close to zero; that is, being of the order of magnitude -2 at worst (with standard deviation being of the order of magnitude -4 at worst). And the average R^2^ for the mean was equal to 1 (that is, standard deviation being zero) and average R^2^ for the standard deviation was consistently 0.999 (with standard deviation being of the order of magnitude -6 at worst). For higher values of f (i.e., 0.5 and 1), the fit was considerably poor (Supplementary Information, Table 9).

### 2.4. Explaining Numerosity Processing Data

In 2002, two studies [10,11], investigating parietal cortex and prefrontal cortex, respectively, observed the presence of neurons whose firing rate was tuned to a specific numerosity. In a study by Nieder and Miller [12], the activity of neurons from the lateral prefrontal cortex displayed a maximum at a preferred number and declined progressively as the numerical distance from that preferred number was increased, thereby forming numerosity filter function. The tuning curves, which had positive skewness on a linear scale, were plotted on nonlinear scales, namely a power function with an exponent of 0.5, a power function with an exponent of 0.33, and a logarithmic scale. The new distributions along with the original distribution (linear scale) were fitted with a Gaussian distribution. The mean goodness-of-fit (R^2^) values were in ascending order when arranged from left to right for the linear scale, the power function with an exponent of 0.5, the power function with an exponent of 0.33, and the logarithmic scale. Additionally, on the logarithmic scale, the variance of numerosity filters was approximately constant with increasing preferred numerosity when the amplitude was kept constant.

As figure 10 shows, output Z_eq_ of C1-FFL is a monotonic function, and hence receptors based on C1-FFL cannot explain these experimental findings. FNM receptors considered in the present study so far produce the maximum output at values which are powers of ten (for example, receptors which constitute the series represented by eqs. 24/25 generate the maximum output at the values 10^1^, 10^2^, 10^3^, and so on up to 10^N^, respectively). Thus, in order to have the receptors produce the maximum output at integral values, such as 1, 2, 3, etc, (Figure 13) and also possess a certain mathematical property, a few combinations of the equilibrium dissociation constants and the Hill-coefficients were tried. The mathematical property that the expression for output Z_eq_ of the individual FNM receptors needed to possess is f_N+1_(X) = f_N_(X/C_N_) when β_1_/α_1_ is kept constant. f_N_ and f_N+1_ are the expressions for output Z_eq_ of consecutive receptors, respectively, and C_N_ is a function of N in general but may also be constant. This property makes the output of the individual receptors with constant β_1_/α_1_ have equal variance when plotted on a logarithm scale, as can readily be seen.

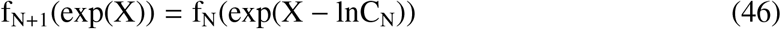

**Figure 13:**
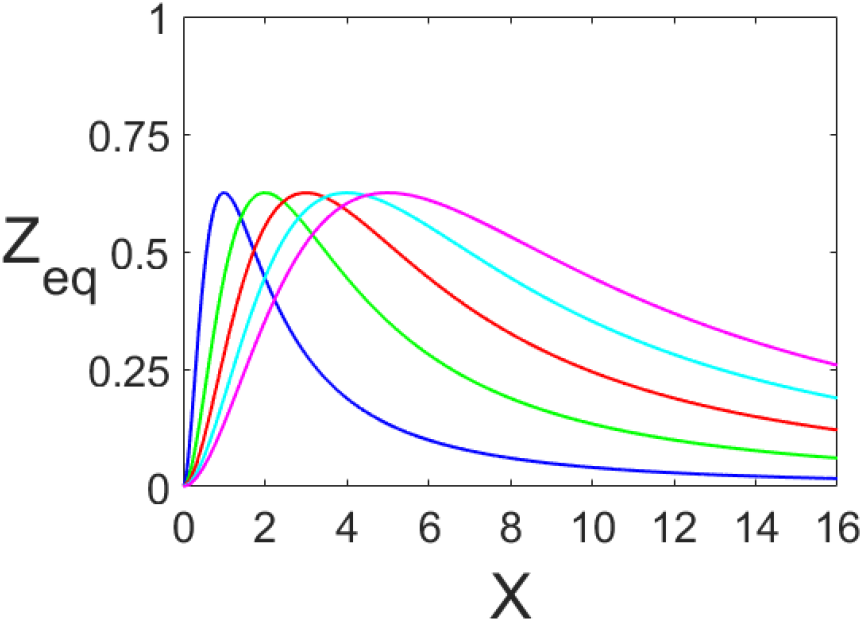
Output Z_eq_ of the receptors which produce the maximum output at integral values. The expression 47 with constant β_1_/α_1_ for N=1,2,3,4,5, respectively, is plotted.

The function f_N+1_ can be got from the function f_N_ by the transformation X → X-lnC_N_ when β_1_/α_1_ is kept constant; hence the standard deviation remains the same. The following function turned out to satisfy both requirements

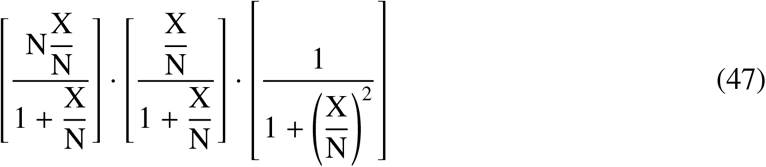

Note that for this function β_1_/α_1_ is equal to N and C_N_ is equal to (N+1)/N.

Like in the study [12], R^2^ values of the Gaussian fit of the expression 47 having constant β_1_/α_1_ (i.e., set equal to 1) for N=1,2,3 plotted on the linear scale, the power function with an exponent of 0.5, the power function with an exponent of 0.33, and the logarithmic scale were in ascending order when arranged from left to right (See Methods, part 5, and Supplementary Information, Table 5).

Notably, barring the last receptor (which does not possess the negative arm), FNM receptors of the series which gives quasi-linear input-output relationship are related in the following way when β_1_/α_1_ is kept constant: f_N+1_(X) = f_N_(X/C_N_), where f is the output function with β_1_/α_1_ set equal to one and N is the receptor’s number. C_N_ for this function is independent of N and is equal to 10. Thus, because output Z_eq_ of the FNM receptor whose series produces quasi-linear input-output relationship follows the above-stated mathematical property when β_1_/α_1_ is kept constant, the output has constant variance when plotted on a logarithm scale. R^2^ values of the Gaussian fit of the first, second, and third term on the RHS of eq. 22 having constant β_1_/α_1_ (i.e., set equal to 1) plotted on the linear scale, the power function with an exponent of 0.5, the power function with an exponent of 0.33, and the logarithmic scale were in ascending order when arranged from left to right (See Methods, part 5, and Supplementary Information, Table 6).

In their experiments, monkeys were shown a set of dots, and then, after a short delay, were presented with another set of dots, which they had to distinguish from the first set. The percentage of trials on which they judged the second number to be indistinguishable from the first number, which varied between 2-6, was plotted as a function of the second number. Monkeys’ performance –percentage of trials on which they reported the two sets to be of the same numerosity– declined progressively as the two sets became numerically more distant. For larger numerosities, the two sets had to be proportionally larger in order to keep the level of performance the same, thereby translating to Weber’ law. The distributions of both monkey’s performance had positive skewness, but assumed a more symmetric shape when plotted on nonlinear scales, namely a power function with an exponent of 0.5, a power function with an exponent of 0.33, and a logarithmic scale. The mean goodness-of-fit values for the linear scale, the power function with an exponent of 0.5, the power function with an exponent of 0.33, and the logarithmic scale were 0.93, 0.97, 0.98, and 0.98, respectively. Further, the variance of the distributions for each numerosity was constant when the data was plotted on a power function scale with 0.33 exponent and the logarithmic scale.

C1-FFL based receptors could not explain the neuronal data, hence they cannot be considered for the possibility of explaining the behavioural data. Because the above-stated formulation of Weber’s law requires the input-output relationship to be (quasi-)linear, the series of FNM receptors which produce the maximum output at integral values must be able to produce (quasi-)linear input-output relationship in order to explain behavioural data of the study by Nieder and Miller [12]. To this end, a series of ten receptors (with the first receptor lacking the positive arm), defined by the expression 47, whose N ranged from 1-10 was considered. The values of β_1_/α_1_ of this series were determined by linear regression (See Methods, part 6). It turned out that the values of β_1_/α_1_ varied over 7 orders of magnitude (R^2^=0.9999), which does not seem biologically plausible. When the series of twenty such receptors (again, with the first receptor lacking the positive arm) were considered, the values of β_1_/α_1_ varied over 6 orders of magnitude (R^2^=1.0000). Thus, the present work is unable to explain the behavioural data.

### 2.5. The Four-node Motif and a Class of Bow-Tie Architecture

As shown in the section *Quasi-linear Input-Output Relationship Over Arbitrary Range*, for the series (of FNM) which generates quasi-linear input-output relationship (eq. 24/25 and figure 4B), b-a+11 receptors are required to provide quasi-linear input-output relationship over the range 10^a^-10^b^. This implies that when a is non-negative and b-a is not too small, 10^b^-10^a^ ≫ b-a+11; and consequently, the generation of quasi-linear relationship between input signal and decoded output becomes an example of bow-tie architecture. This gets more pronounced as both b and a becomes large. For example, when a=0 and b=5, the range of the input signal and decoded output (10^5^-1) is four orders of magnitude larger than the number of receptors (16). Similarly, as shown in the section *The Mathematical Basis of the Quasi-Linear Input-Output Relationship Provided by the Series of the Four-node Motif*, for the series (of C1-FFL) which generates linear input-output relationship (eq. 28/29 and figure 8B), b-a+9 receptors are required to provide linear input-output relationship over the range 10^a^-10^b^. Again, this implies that when a is non-negative and b-a is not too small, 10^b^-10^a^ ≫ b-a+9; and consequently, the generation of linear relationship between input signal and decoded output becomes an example of bow-tie architecture. This gets more pronounced as both b and a becomes large. For example, when a=0 and b=5, the range of the input signal and decoded output (10^5^-1) is four orders of magnitude larger than the number of receptors (14). Therefore, a class of bow-tie architecture can be defined: a class in which the number of receptors is much smaller than the range of input signal and decoded output.

The range of output Z_eq_ generated by the series of FNM (eq. 24/25 and figure 4B) and the series of C1-FFL (eq. 28/29 and figure 8B) are identical to the input signal’s range. In contrast to that, now I consider a hypothetical requirement of sensitive detection over wide range of input signal when the output has an upper limit. Black line in figure 14A depicts that when the output has an upper limit, sensitivity of a mechanism generating single linear input-output relationship reduces as the input signal’s range increases. But the sensitivity is maintained if a mechanism generates multiple linear input-output relationships, each of which is from a different receptor and covers corresponding limited range of input signal. The blue, green, and red lines are ideal hypothetical outputs which together provide sensitive detection over a wide range of input signal when the output has an upper limit. However, the outputs are not biologically realistic, as they terminate abruptly at their maximum value. Figure 14B shows a biologically plausible form of the outputs.

**Figure 14:**
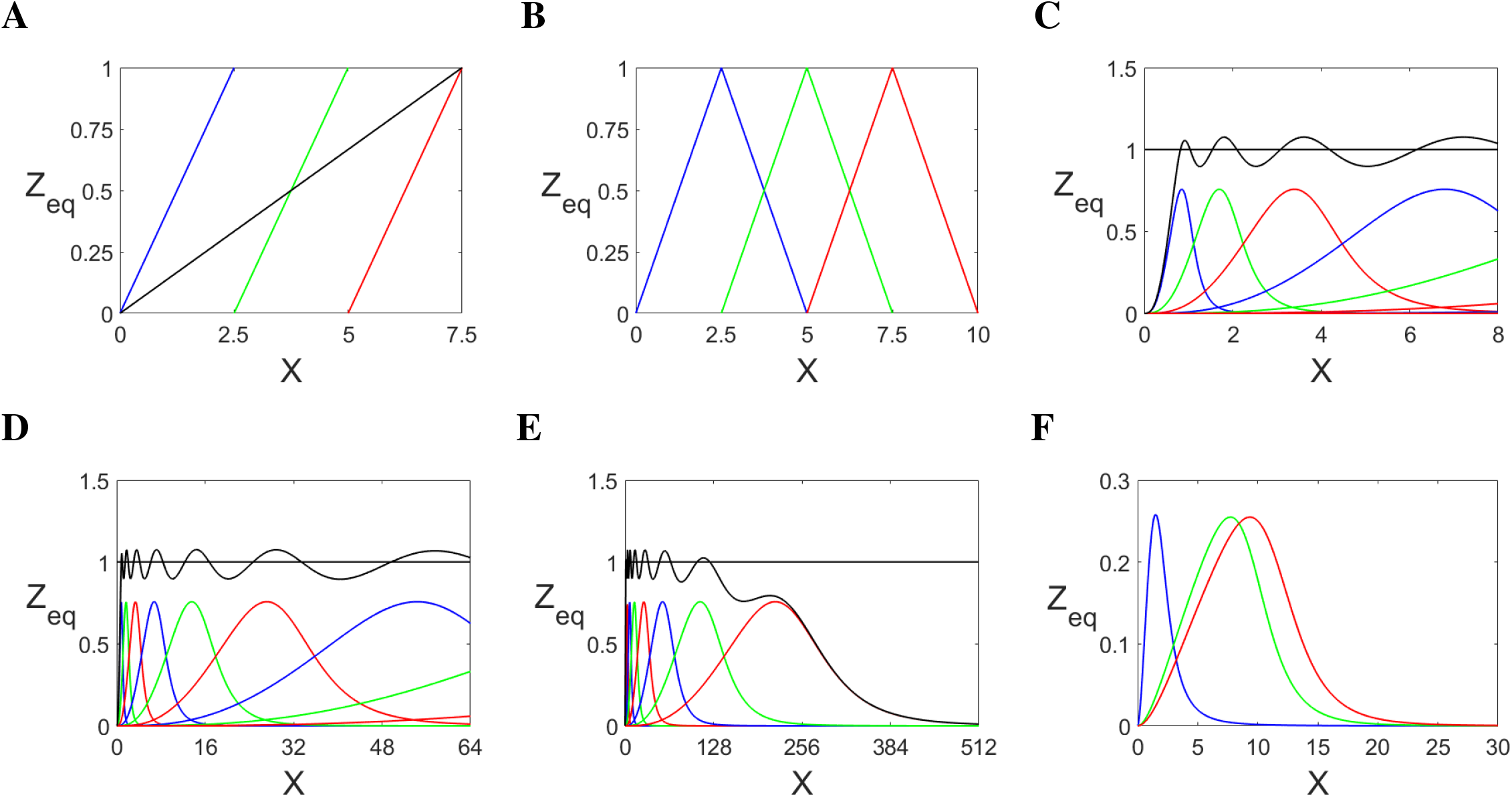
(**A**) The black line shows that, when the output has an upper limit, sensitivity of linear input-output relationship reduces with increase in the range of input signal. The blue, green, and red lines are minimal ideal hypothetical input-output relationship, i.e., which maintain sensitivity throughout the input signal’s range. (**B**) Biologically realistic form of the ideal hypothetical input-output relationship. (**C-E**) Output Z_eq_ of the series of receptors which are defined by the expression 48 (with K=5) for various ranges of N. The individual outputs are approximately Gaussian and have constant coefficient of variation. (**F**) A series of three FNM receptors is able to qualitatively generate the absorption spectra of human cone cells. The expressions of the three receptors are: [3.6 · (X/4)/(1 + X/4)] · [(X/8)^2^/(1 + (X/8)^2^)] · [1/(1 + (X/8)^8^)]; [5 · (X/20)/(1 + X/20)] · [(X/20)^2^/(1 + (X/20)^2^)] · [1/(1 + (X/20)^8^)]; [5 · (X/24)/(1 + X/24)] · [(X/24)^2^/(1 + (X/24)^2^)] · [1/(1 + (X/24)^8^)]

Figure 14C shows output Z_eq_ of a series of FNM which qualitatively matches figure 14B and, thus, provides sensitive detection over arbitrary range of input signal when the output has an upper limit. FNM in this series has the following properties: β_1_/α_1_ (K) is constant, the Hill-coefficients m and n are equal to 2 and 8, respectively, all equilibrium dissociation constants of a given receptor are equal, and the value of equilibrium dissociation constants of a receptor are twice those of the preceding receptor. The expression 48 defines the general form of this receptor. Z_eq_ of this FNM fits Gaussian distribution very well (See Methods, part 7). R^2^ for the Gaussian fit of the expression 48 for N=1,2,3,4,5 are 0.9965, 0.9955, 0.9954, 0.9953, and 0.9953, respectively. Although the output when m=2, n=10 and m=1, m=8, respectively, for equilibrium dissociation constants equal to 1, give slightly better R^2^ (Supplementary Information, Table 7), they are visibly less symmetric than the output when m=2, n=8 (Figure 15).

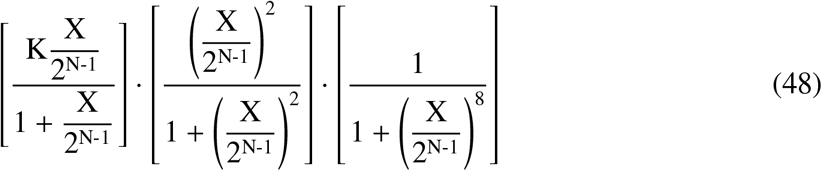

**Figure 15:**
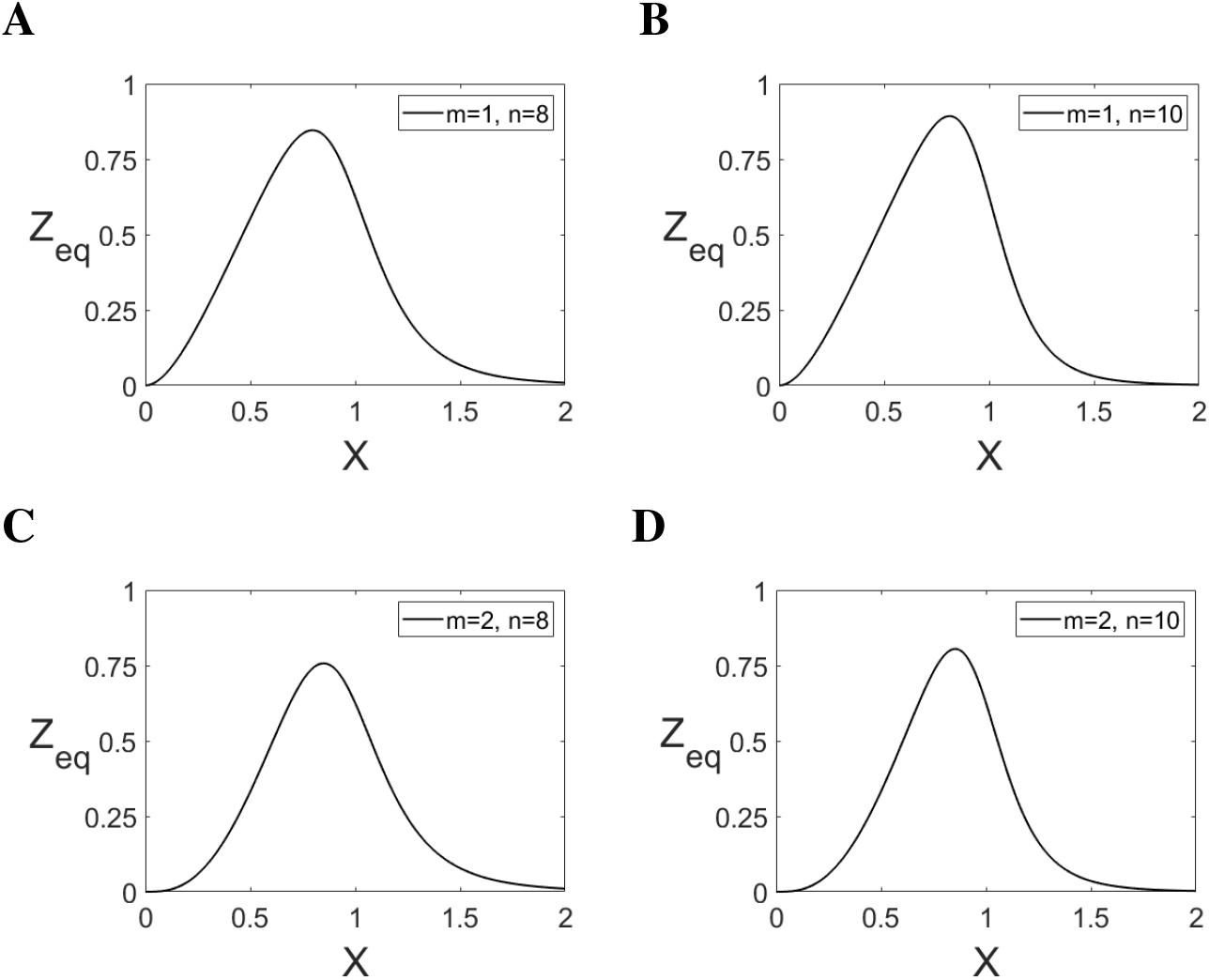
(**A-D**) Output Z_eq_ of FNM defined by the expression [5 · (X/1)/(1 + X/1)] · [(X/1)^m^/(1 + (X/1)^m^)] · [1/(1 + (X/1)^n^)] with different values of the Hill-coefficients m and n.

Because the expression 48, whose β_1_/α_1_ is already a constant, possesses the mathematical property f_N+1_(X) = f_N_(X/C_N_), where C_N_ is independent of N and is equal to 2, the coefficient of variation of Gaussians they are approximately equivalent to is constant. That is, the mean and standard deviation of the output of a receptor is C_N_ times the mean and standard deviation of the output of the preceding receptor. Further, between the mean of the output of the first receptor and the antepenultimate receptor, the total output of the series of this receptor oscillates with constant amplitude and decreasing frequency around a flat line. Note that the individual outputs in figure 14B have constant width at half height (WAHH); whereas, because WAHH of Gaussian distribution is equal to 4·ln2·σ, WAHH of the outputs in figure 14C are proportional to the mean of the approximate Gaussians they are equivalent to. It should be noted that even though the output’s range has an upper limit, the range of the decoded output is still equal to the input signal’s range, just like for the series of FNM (eq. 24/25) and C1-FFL (eq. 28/29) which generate (quasi-)linear input-output relationship. That’s because for a given input signal, the total output is a unique combination of the output of the individual receptors. Thus, for a given input signal, a unique output can be decoded from the output of the individual receptors. Thus, for the series of receptors that are defined by the expression 48, the decoded output is different from output Z_eq_, whereas for the series defined by eqn. 24/25 and 28/29, the two outputs are equal. This implies that the biochemical nature of the output Z_eq_ of all receptors of the series defined by eqn. 24/25 and 28/29 should be identical, whereas the biochemical nature of the output Z_eq_ of receptors of the series whose terms are defined by the expression 48 should be specific for every receptor. That is, in the latter series, Z_eq_ of a receptor decodes to a higher value than Z_eq_ of the receptor that precedes it.

Assuming the equilibrium dissociation constant of the first and the last receptor to be 2^a^ and 2^b^, respectively; arguably, the range in which an input signal can be detected sensitively is µ_1_·(2^b^-2^a^) + (1/2) · σ_1_·(2^a^+2^b^), where b > a and both are integers, and µ_1_ (= 0.8461) and σ_1_ (= 0.2748) are the mean and standard deviation of the Gaussian distribution which fits output Z_eq_ of the receptor whose equilibrium dissociation constants are equal to 1 (i.e., K=5 and N=1 in the expression 48). The number of receptors in this series is equal to b-a+1. This implies that when a is non-negative and b-a is not too small, µ_1_·(2^b^-2^a^) + (1/2) · σ_1_·(2^a^+2^b^) ≫ b-a+1; and consequently, the linear relationship between input signal and decoded output becomes an example of bow-tie architecture. This gets more pronounced as both b and a becomes large. For example, when a=0 and b=10, the range of the input signal and decoded output (1006.4) is two orders of magnitude larger than the number of receptors (11). Figure 14D shows that, with an intuitive choice of parameters, a series of three such FNM receptors is able to produce an input-output relationship which qualitatively matches absorption spectra of three cone opsins of humans [13]. The defining expression of the three receptors are in the figure’s legend. This implies that FNM is likely to be involved in the perception of the visible light spectrum in humans and presumably in other organisms as well. Note that this absorption spectra is a bow-tie architecture: three cone opsins perceive a portion of the electromagnetic spectrum. Because in many organisms sound frequency perception occurs over several orders of magnitude [14], thereby forming a bow-tie architecture, FNM might be involved in the perception of sound frequency also.

### 2.6. The Four-node Motif and Logarithm Computation

When the ratio β_1_/α_1_ in the series of FNM defined by eq. 24 increases as powers of ten, Z_eq_(X) is a straight line. And when the ratio β_1_/α_1_ increases linearly, the output of the same series is a concave curve (Figure 16A). Thus it was suspected that with rational choice of β_1_/α_1_, the output could be equal to the logarithm of the input signal. Because a series was sought which could compute logarithm of input signal ranging between 1 and 10^4^, series of five FNM receptors (eq. 24) having Kd_XZ_ equal to 10^1^, 10^2^, 10^3^, 10^4^, and 10^5^, respectively, was considered. The values of β_1_/α_1_ of the series which compute logarithm with the base ten and the natural logarithm were estimated by linear regression (See Methods, part 8). It turned out that the series is able to compute logarithm of the input signal (Figure 17,18). The same procedure was carried out on the series of C1-FFL (eq. 28), as its output also is a concave curve when the ratio β_1_/α_1_ increases linearly (Figure 16B). It turned out that this series also is able to compute logarithm of the input signal (Figure 19,20). The computation of logarithm by the series of FNM and C1-FFL raised the possibility that, with rational choice of β_1_/α_1_, the two series might also be able to compute the exponential of the input signal. As before, linear regression was used to find out the values of β_1_/α_1_; however, none of the series was able to generate the exponential of the input signal (See Methods, part 8).

**Figure 16:**
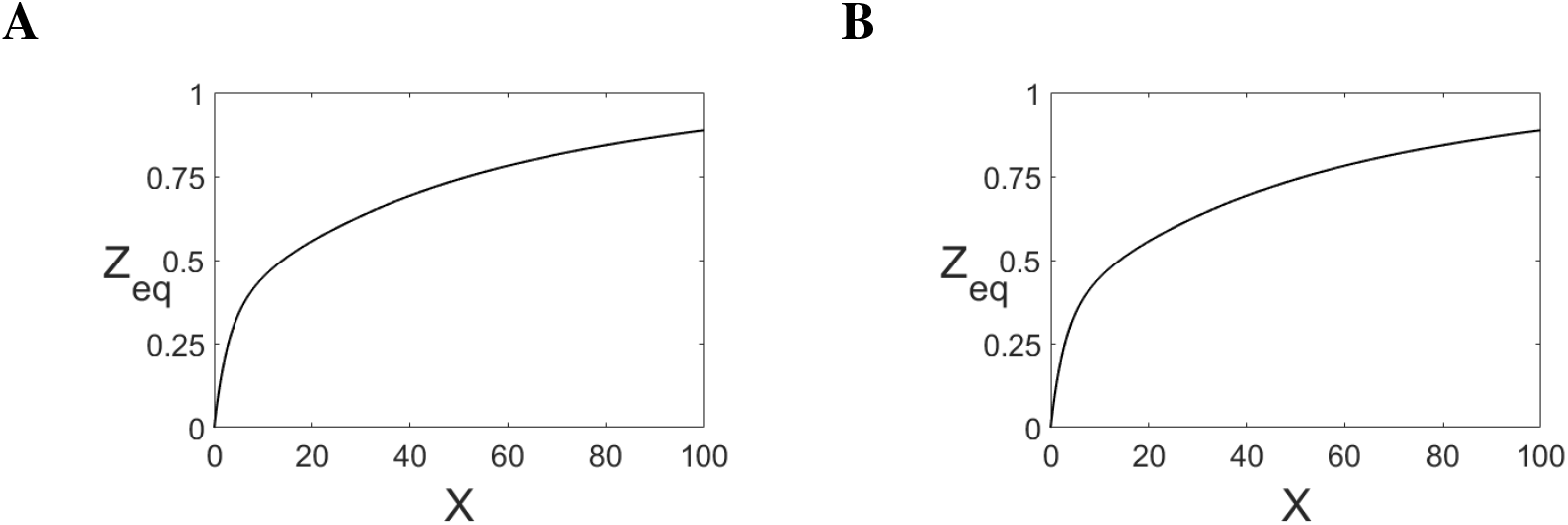
Output Z_eq_ of the series of FNM (**A**) and C1-FFL (**B**), respectively, when β_1_/α_1_ of the five receptors are 1,2,3,4,5, respectively.

**Figure 17:**
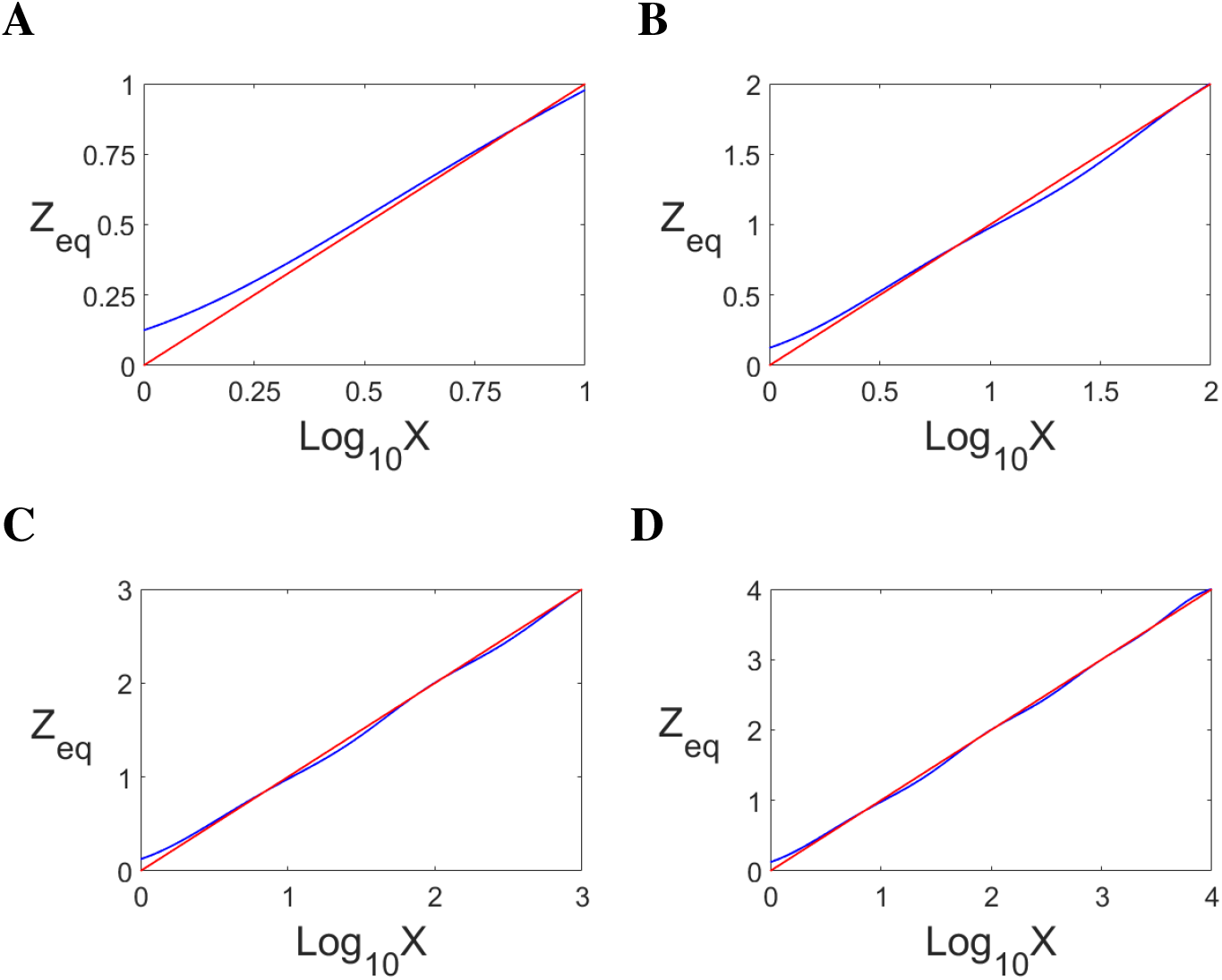
(**A-D**) Output of the series of FNM which computes log_10_. The total output (blue) of the first five receptors for various range of input signal plotted on log_10_-transformed abscissa. The straight line (red), which is the logarithm function plotted on a log-transformed abscissa, is drawn for comparison. The first receptor possesses the positive arm, with m=2, in order to dampen the output when the input signal is below one. The last receptor lacks the negative arm. β_1_/α_1_ of the five receptors were: 2.89, 6.299, 8.587, 12.36, 8.317, respectively. The R^2^ was 0.9989.

**Figure 18:**
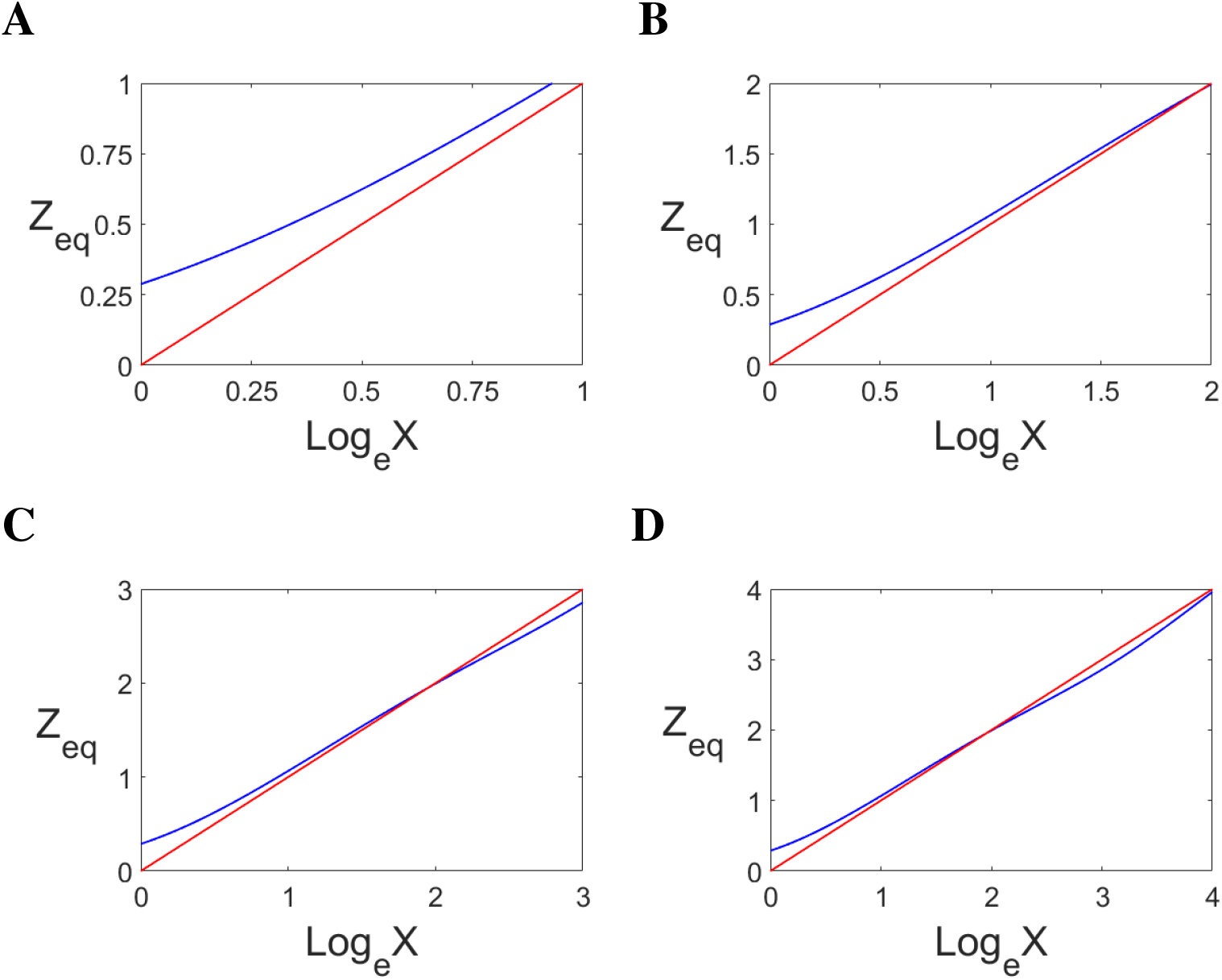
(**A-D**) Output of the series of FNM which computes log_e_. The total output (blue) of the first five receptors for various range of input signal plotted on log_e_-transformed abscissa. As previously, the value of m of the first receptor’s positive arm is set equal to 2 and the last receptor lacks the negative arm. β_1_/α_1_ of the five receptors were: 6.654, 14.5, 19.77, 28.47, 19.15, respectively. The R^2^ was 0.9989.

**Figure 19:**
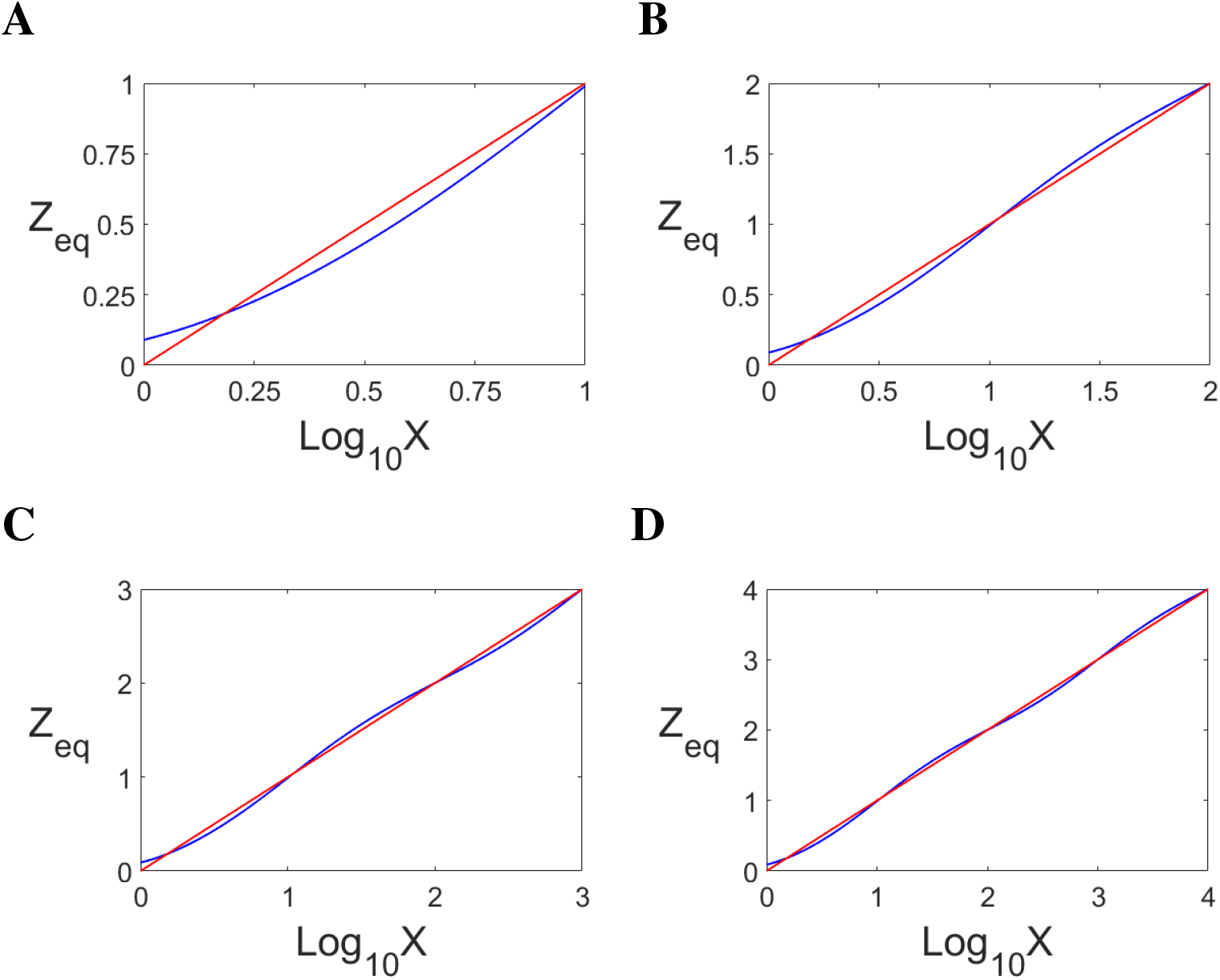
(**A-D**) Output of the series of C1-FFL which computes log_10_. The total output (blue) of the first five receptors for various range of signal plotted on log_10_-transformed abscissa. The straight line (red), which is the logarithm function plotted on a log-transformed abscissa, is drawn for comparison. The first receptor possesses the positive arm, with m=2, in order to dampen the output when the input signal is below one. β_1_/α_1_ of the five receptors were: 1.97 0.292 1.697 0.3015 1.736, respectively. The R^2^ was 0.9995.

**Figure 20:**
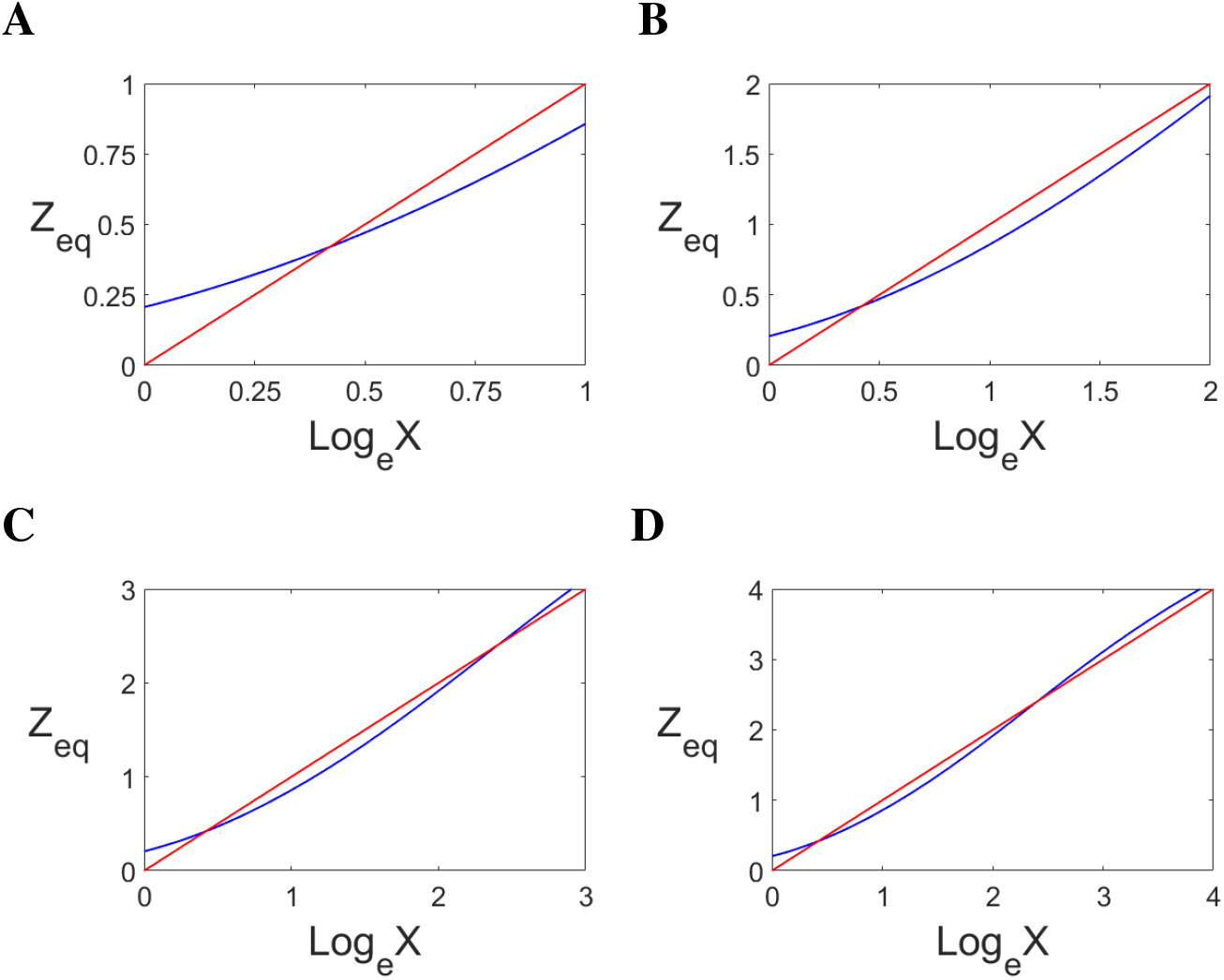
(**A-D**) Output of the series of C1-FFL which computes log_e_. The total output (blue) of the first five receptors for various range of signal plotted on log_e_-transformed abscissa. As previously, the value of m of the first receptor’s positive arm is set equal to 2. β_1_/α_1_ of the five receptors were: 4.536 0.6723 3.907 0.6941 3.996, respectively. The R^2^ was 0.9995.

Table 3 and 4 depict how the parameters of different series based on FNM and C1-FFL scale with the receptor’s number, respectively. The value of (1/Kd_YZ_) · (β_2_/α_2_) and (1/Kd_VZ_) · (β_3_/α_3_) for the series based on FNM and C1-FFL is equal to 10^4^ and 10^8^, respectively. Those equilibrium dissociation constants whose individual values are not known (i.e, which are lumped together with other parameters) are marked NA.

**Table 3:** Type of Series based on FNM and the Values of its Parameters

| Series Type | $\beta_1/\alpha_1$ | $Kd_{XZ}$ | $Kd_{XY}$ | $Kd_{XV}$ | m | n |
| --- | --- | --- | --- | --- | --- | --- |
| Linear input-output | $Kd_{XZ}$ | Powers of 10 | $10^4 \cdot Kd_{XZ}$ | $10^3 \cdot Kd_{XZ}$ | 1 | 1 |
| Numerosity Processing in Monkey | $Kd_{XZ}$ | Integral values | NA | NA | 1 | 2 |
| Sensitive Detection when the Output has an Upper Limit | Constant | Powers of 2 | NA | NA | 2 | 8 |
| Logarithm Computation | Determined by Linear Regression | Powers of 10 | $10^4 \cdot Kd_{XZ}$ | $10^3 \cdot Kd_{XZ}$ | 1 | 1 |

**Table 4:** Type of Series based on C1-FFL and the Values of its Parameters

| Series Type | $\beta_1/\alpha_1$ | $Kd_{XZ}$ | $Kd_{XY}$ | $Kd_{XV}$ | m | n |
| --- | --- | --- | --- | --- | --- | --- |
| Linear input-output | $Kd_{XZ}$ | Powers of 10 | $10^8 \cdot Kd_{XZ}$ | $10^7 \cdot Kd_{XZ}$ | 1 | 1 |
| Logarithm Computation | Determined by Linear Regression | Powers of 10 | $10^8 \cdot Kd_{XZ}$ | $10^7 \cdot Kd_{XZ}$ | 1 | 1 |

## 3. Methods

Part 1a

The rationale behind setting (1/Kd_YZ_)·(β_2_/α_2_) and (1/Kd_VZ_) · (β_3_/α_3_)) for FNM equal to 10^4^ is as follows.

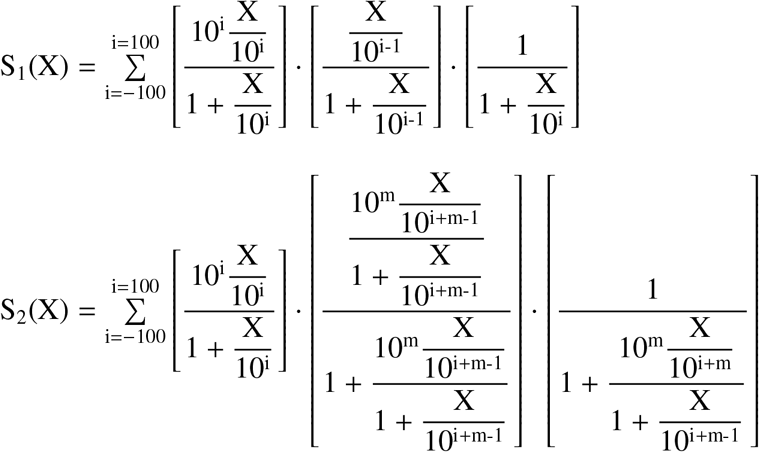

That value of the two quantities were chosen for which 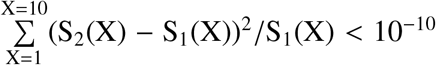. The minimum value of m turned out to be 4.

Part 1b

The rationale behind setting (1/Kd_YZ_)·(β_2_/α_2_) and (1/Kd_VZ_) · (β_3_/α_3_)) for C1-FFL equal to 10^8^ is as follows.

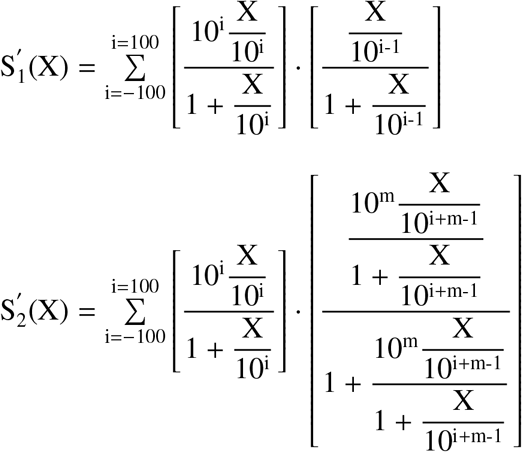

That value of the two quantities were chosen for which 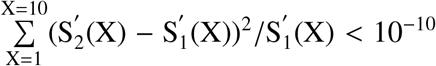. The minimum value of m turned out to be 8.

Part 2a

How well the first term on the RHS of eq. 22 approximates the first term on the RHS of eq. 23 was determined by regressing the output of the first term on the RHS of eq. 23 against the first term on the RHS of eq. 22 multiplied by a coefficient (RHS of eq. 49). Note that the first term on the RHS of eq. 22/23 defines the first receptor of the series of FNM which provides quasi-linear input-output relationship. As stated above, this receptor is meant to perceive the input signal (X) whose value is one order of magnitude less than Kd_XZ_ of the receptor. Also, because Kd_XZ_ of the first receptor is equal to 10, its output peaks when X is equal to 10. Thus, for the purpose of (linear) regression, the output of the first term on the RHS of eq. 23 was evaluated up to one order of magnitude more than the Kd_XZ_; that is, at the following values of X: 1, 2, 5, 10, 20, 50, 100. At X equal to 100, the output has declined to 36% of its peak value.

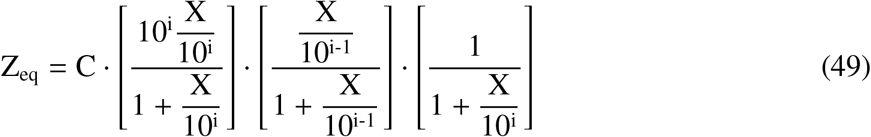

If the approximation is good, the value of the coefficient should turn out to be around one, besides goodness-of-fit (r^2^) being close to one. The value of the coefficient for all the combinations of m and n was close to one. The goodness-of-fit was equal to one for all the combinations (Supplementary Information, Tables 1-3). Specifically, for m and n equal to one, the value of C was 1.0000.

Part 2b

How well the first term on the RHS of eq. 28 approximates the first term on the RHS of eq. 29 was determined by regressing the output of the first term on the RHS of eq. 29 against the first term on the RHS of eq. 28 multiplied by a coefficient (RHS of eq. 50). As previously, for the purpose of (linear) regression, the output of the first term on the RHS of eq. 29 was evaluated up to one order of magnitude more than the Kd_XZ_; that is, at the following values of X: 1, 2, 5, 10, 20, 50, 100. At X equal to 100, the output is 90% of its maximum value (i.e., the asymptotic output at a very large X).

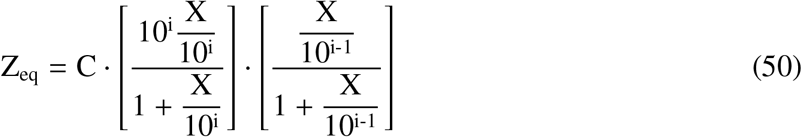

If the approximation is good, the value of the coefficient should turn out to be around one, besides goodness-of-fit (r^2^) being close to one. The value of the coefficient for every m was equal to one. The goodness-of-fit was equal to one for every m (Supplementary Information, Table 4).

Part 3

Fitting straight line with output Z_eq_ given by eq. 25 (the non-approximated form) for various ranges of Kd_XZ_ and input signal (Table 1). For a given range of input signal, the output was generated at the input values which were powers of 10. For example, for the input signal’s range 10^−1^–10^3^, the output was generated at the following values: 10^−1^, 10^0^, 10^1^, 10^2^, and 10^3^. The same procedure was carried out after shifting the range of Kd_XZ_ and input signal, shown in table 1, by the factor of 0.01, 0.1, 10, and 100. That is, for example, for the second case (factor=0.1) the new ranges of Kd_XZ_ were 10^−3^–10^3^, 10^−4^–10^4^, 10^−5^–10^5^, 10^−6^–10^6^, and 10^−7^–10^7^. And the new ranges of input signal were 10^−1^–10^1^, 10^−2^–10^2^, 10^−3^–10^3^, 10^−4^–10^4^, and 10^−5^–10^5^. Similarly, for the third case (factor=10) the new ranges of Kd_XZ_ were 10^−1^–10^5^, 10^−2^–10^6^, 10^−3^–10^7^, 10^−4^–10^8^, and 10^−5^–10^9^. And the new ranges of input signal were 10^1^–10^3^, 10^0^–10^4^, 10^−1^–10^5^, 10^−2^–10^6^, and 10^−3^–10^7^. The same procedure was also carried out with output Z_eq_ given by eq. 28 (the non-approximated form) for various ranges of Kd_XZ_ and input signal (Table 2), in order to determine if the series of C1-FFL could generate linear input-output relationship.

Part 4

The formalism of Weber’s law presented in this paper; that is, the output Z_eq_ given by eq. 25 (approximated as eq. 24) and eq. 29 (approximated as eq. 28) is a random sample from the Gaussian whose mean is equal to the input signal and standard deviation is equal to the Weber fraction times the mean, is validated as follows. As stated above, in order to validate this assumption, output Z_eq_ of eq. 25 and eq. 29, respectively, should be shown to be a Gaussian distribution with mean equal to a constant times the input signal’s value and standard deviation equal to a constant times the Weber fraction times the mean when the output generated by individual components of FNM and C1-FFL, respectively, is a random sample from their corresponding Gaussian distributions. That is, given the input signal X (taken as multiple of 10), the value of V and Y are generated from Gaussian distribution with mean equal to 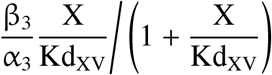 and 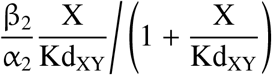 respectively, and standard deviation equal to the Weber fraction times the respective mean.

Then, with the value of X, V and Y, Z_eq_ is generated from Gaussian with mean equal to

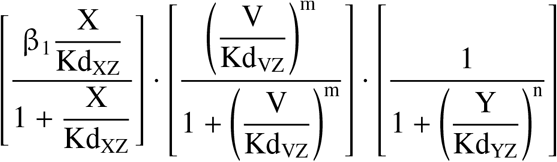

and standard deviation equal to the Weber fraction times the mean. Adding output Z_eq_ of all receptors of the series and multiplying it by the reciprocal of 0.7489 (the limiting value of the slope when the input signal is a multiple of 10) and 1.111 for the series of FNM and C1-FFL, respectively, should turn out to be equivalent to sampling from a Gaussian with mean equal to a constant times the input signal X and standard deviation equal to a constant times the Weber fraction times the input signal X. As stated above, in order to get (quasi-)linear input-output relationship over the range 10^a^-10^b^ (where, a < b and both are integers) from the series of FNM and C1-FFL, Kd_XZ_ of the receptors should range at least from 10^a-5^ to 10^b+5^ and 10^a-4^ to 10^b+4^, respectively. Thus, the input-output relationship is (quasi-)linear only for the middle values of input signal, and therefore the order of magnitude of X should be among the preferred order of magnitude of the middle receptors.

The value of (β_2_/α_2_) · (1/Kd_YZ_) and (β_3_/α_3_) · (1/Kd_VZ_) were set equal to 10^4^ (eqs. 9 and 20) and 10^8^ for FNM and C1-FFL, respectively, in order to carry out the approximations. However, values of β_2_/α_2_ and β_3_/α_3_, required to generate the value of V and Y from the input signal X, are not known individually. Therefore three combinations of β_2_/α_2_ and Kd_YZ_ were considered, while keeping the value of the product of β_2_/α_2_ and 1/Kd_YZ_ equal to 10^4^ and 10^8^ for FNM and C1-FFL, respectively. The combinations for FNM were β_2_/α_2_=10^4^ and Kd_YZ_=1, β_2_/α_2_=10^2^ and Kd_YZ_=10^−2^, and β_2_/α_2_=1, Kd_YZ_=10^−4^. The combinations for C1-FFL were β_2_/α_2_=10^8^ and Kd_YZ_=1, β_2_/α_2_=10^4^ and Kd_YZ_=10^−4^, and β_2_/α_2_=1, Kd_YZ_=10^−8^. The same was done with β_3_/α_3_ and Kd_VZ_ for FNM and C1-FFL.

Given a value of the mean (equivalent to the input signal X) and the Weber fraction (f), ten thousand values of Z_eq_ were generated from this sampling procedure and used to fit a Gaussian distribution. This procedure was carried out for thirty times. For a given value of f, the average of (thirty) means of the Gaussian fit for mean (the input signal X) equal to 0.1, 1, and 10 (and therefore, standard deviation equal to f·0.1, f·1, and f·10, respectively) were linearly regressed against the following values: 0.1, 1, and 10. Similarly, for a given value of f, the average of (thirty) standard deviations got from the same procedure were linearly regressed against the following values: f·0.1, f·1, and f·10. The values of f taken are as follows: 0.01, 0.02, 0.05, 0.1, 0.2, 0.5, and 1. This was done for all the nine combinations of β_2_/(α_2_ · Kd_YZ_) and β_3_/α_3_·Kd_VZ_). Table 8 and 9 of Supplementary Information show the average and standard deviation of the slope, y-intercept, and R^2^ of the straight line fitting of the average of thirty means and standard deviations (derived from Gaussian fitting of Z_eq_) for the series of FNM and C1-FFL, respectively. The value of the average was truncated after two decimal places, and the value of standard deviation was truncated after one decimal place.

Part 5

For the purpose of Gaussian fitting, the expression 47 for N=1,2,3, plotted on the linear scale, the power function with an exponent of 0.5, the power function with an exponent of 0.33, and the logarithmic scale, were evaluated at values of X which were uniformly spaced with the interval of 0.25 and ranged from 0 to the value at which the output had declined to 0.01 times the peak value. The same was done with the first three terms on the RHS of eq. 22, except for the linear scale, for which the range taken was from 0 to the value at which the output had declined to 0.5 times the peak value. This was done in order to reduce the number of data points towards the right side of the peak. For both fittings (except for the special case mentioned), reducing the interval further caused a minuscule change in the R^2^ but did not change the result qualitatively (Supplementary Information, Table 5 and 6, respectively).

Part 6

In order to get the series of FNM receptors which produce the maximum output at integral values generate (quasi-)linear input-output relationship, the values of β_1_/α_1_ of this series were determined by linearly regressing output Z_eq_ of the series of ten such receptors, defined by the expression 47 with N ranging from 1-10, on the values of X which were uniformly spaced with the interval of 0.25 and ranged from 0.25 to 10. The same was done with the series of twenty such receptors, defined by the expression 47 with N ranging from 1-20. The positive arm of the first receptor in both the series was removed, as its presence gave an uneven input-output curve mostly towards its leftmost part.

Part 7

For the purpose of Gaussian fitting, the expression 48, with different combinations of m and n, was evaluated at values of X which were uniformly spaced with the interval of 0.25 and ranged from 0 to the value at which the output had declined to 0.01/K times the peak value. K was set equal to 5. Reducing the interval further caused a minuscule change in the R^2^ but did not change the result qualitatively (Supplementary Information, Table 7).

Part 8

In order to determine β_1_/α_1_ for the computation of the logarithm by the series of FNM and C1-FFL, the output of log_10_(X) and log_e_ (X) were linearly regressed against the expression on the RHS of eq. 51 and eq. 52, respectively. To this end, the functions log_10_(X) and log_e_ (X) and the expressions were evaluated at values which were powers of 10, ranging from 10^0^ to 10^6^ . For the range 10^0^ to 10^5^, the value of three out of five β_1_/α_1_ turned out to be negative for both series. Note that at least five input values were needed for the regression procedure, as the series of five receptors was considered. Because the logarithm of values that lie between 0 and 1 is negative, a biologically unrealistic scenario, the output in this range was suppressed. This was done by setting the Hill-coefficient of the positive arm of the first receptor equal to two (i.e., m=2). For the series of C1-FFL, it was observed that the first receptor with m=1 produced two negative β_1_/α_1_. If the maximum value of the range was more than 10^6^ (at least up to 10^15^), the value of all β_1_/α_1_ remained positive for the series of FNM. However, the fitting procedure returned at least one negative β_1_/α_1_ for the series of C1-FFL when the maximum value of the range was more than 10^6^ (at least up to 10^15^). These two observations suggest that the series of FNM is more robust in computing logarithm than the series of C1-FFL. In order to determine β_1_/α_1_ for the computation of exponential by the series of FNM and C1-FFL, the output of exp(X) was linearly regressed against the expression on the RHS of eq. 51 and eq. 52, respectively. To this end, the function exp(X) and the expressions were evaluated at values which were square root of numbers ranging from 0 to 6.

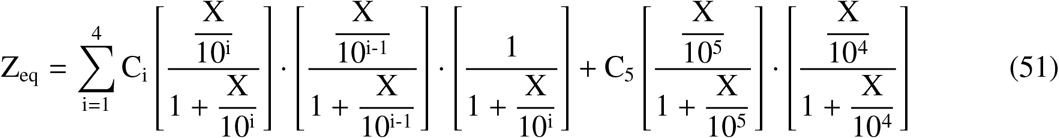

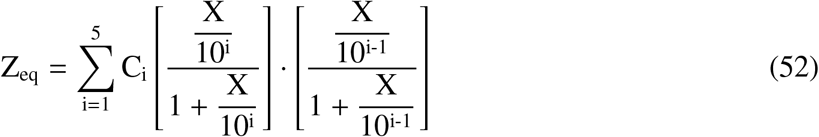

## 4. Discussion

Sensory systems need to respond to stimuli which vary over several orders of magnitude in intensity and frequency. One way to achieve this is by “resetting” the system, that is, by employing the mechanism of exact adaptation. A prominent example of this strategy is *Escherichia coli*’s chemotaxis. Tu and co-workers [4] presented a three-variable reduced model of the bacterial chemotaxis, whose features are: 1) attractant ligand (L(t)) binding reduces the average kinase activity (a(t)), 2) the kinase activates demethylation machinery (the time rate of change in methylation must be a monotonically decreasing function of the kinase activity), and 3) attractant ligand binding affinity is a decreasing function of the average methylation level (m(t)). Binding of an attractant ligand lowers the average kinase activity (and thus decreases the tumbling frequency), leading to reduction of demethylation rate, and the resultant increased level of methylation restores the average kinase activity back to its resting value.

The sensory mechanism presented in this work in order to explain Weber’s law in higher-order sensory processes differs fundamentally from the exact adaptation based mechanism, which is responsible for Weber’s law observed in E coli’s chemotaxis. The two differences are as follow. Firstly, in the latter mechanism an input signal is sensed relative to the background signal, at which the system was equilibrated to. By contrast, in the former mechanism an input signal is sensed in absolute manner. Secondly, in the latter mechanism it’s the transient response of the system that possesses information about the input signal, as the system’s output returns back to its resting value. On the other hand, in the former mechanism the equilibrium state of the system possesses information about the input signal. Because higher-order sensory processes such as weight perception, size perception, and numerosity processing sense an input signal in absolute manner and should not involve exact adaptation (as argued in *Introduction*), a sensory mechanism based on exact adaptation cannot explain Weber’s law observed in the three higher-order sensory processes. Thus the mechanism proposed here is, to the author’s knowledge, the only plausible mechanism which can explain Weber’s law observed in the three higher-order sensory processes.

Here the present author would like to speculate on the role of numeral systems. Arguably, estimating the number of an entity without using a numeral system amounts to generating an output which is the input signal plus noise which is proportional to the input signal; that is, Weber’s law. Thus, arguably, the function of a numeral system is to remove the noise and have the brain generate an output which is equal to the input signal. However, because large numbers are considered in terms of their order of magnitude; arguably, the brain processes large numbers in accordance with Weber’s law.

It should be noted that there has to be an upper limit on the production rate (β_1_). However, because eq. 2 (and hence also eq. 18/21) is linear in β_1_, the overall output can be increased by having the input signal engender the production from multiple *Z* producers. There also has to be an upper limit on the value of the equilibrium dissociation constants, so that the Gibbs energy of binding remains significantly larger than the thermal energy. Thus, some mechanism has to exist such that the apparent value of the equilibrium dissociation constants increases even when the equilibrium dissociation constants for the actual binding reach an upper limit. This can be achieved, for example, by having a mechanism in which the input signal has to occupy multiple producer elements of the producer *Z* at the same time in order to engender production of Z. An implication of these two considerations is that FNM and C1-FFL might be present in a hidden form in biological networks.

