## Supplementary Information for "The Series of a Four-node Motif and Coherent Type-1 Feed-forward Loop can Provide Sensitive Detection over Arbitrary Range of Input Signal, thereby Explain Weber’s Law in Higher-Order Sensory Processes, and Compute Logarithm"

The next three tables (1-3) show the value of the coefficient C and the goodness of fit ($R^2$) obtained from linearly regressing the output of the first term on the RHS of equation 25 (which defines an FNM) against the RHS of equation 49, for different combinations of the Hill coefficients m and n.

**Table 1:** m is equal to 1, while n varies from 1 to 16

| **Hill Coefficients (across)** | m=1, n=1 | m=1, n=2 | m=1, n=4 | m=1, n=8 | m=1, n=16 |
|---|---|---|---|---|---|
| C | 1.0000 | 1.0000 | 1.000 | 1.0001 | 1.0002 |
| $R^2$ | 1.0000 | 1.0000 | 1.0000 | 1.0000 | 1.0000 |

The table 4 shows the value of the coefficient C and the goodness of fit ($R^2$) obtained from linearly regressing the output of the first term on the RHS of equation 29 (which defines a C1-FFL) against the RHS of equation 50, for different values of the Hill coefficients m.

**Table 2:** m varies from 2 to 16, while n is equal to 1

| **Hill Coefficients (across)** | m=2, n=1 | m=4, n=1 | m=8, n=1 | m=16, n=1 |
|---|---|---|---|---|
| C | 1.0001 | 1.0001 | 1.0001 | 1.0001 |
| $R^2$ | 1.0000 | 1.0000 | 1.0000 | 1.0000 |

**Table 3:** m and n are equal and vary from 2 to 16

| **Hill Coefficients (across)** | m=2, n=2 | m=4, n=4 | m=8, n=8 | m=16, n=16 |
|---|---|---|---|---|
| C | 1.0001 | 1.0001 | 1.0001 | 1.0002 |
| $R^2$ | 1.0000 | 1.0000 | 1.0000 | 1.0000 |

**Table 4:** m is equal to 1, while n varies from 1 to 16

| **Hill Coefficients (across)** | m=1 | m=2 | m=4 | m=8 | m=16 |
|---|---|---|---|---|---|
| C | 1.0000 | 1.0000 | 1.0000 | 1.0000 | 1.0000 |
| $R^2$ | 1.0000 | 1.0000 | 1.0000 | 1.0000 | 1.0000 |

**Table 5:** $R^2$ of the Gaussian Fit of the Expressions 47 for N=1,2,3 Plotted on Linear, Power Function-compressed, and Logarithmically Compressed Axes

| Linear | Power Function (0.5) | Power Function (0.33) | Logarithm |
|---|---|---|---|
| 0.8442 | 0.9328 | 0.9937 | 0.999 |
| 0.8535 | 0.9327 | 0.9917 | 0.9977 |
| 0.8571 | 0.9328 | 0.991 | 0.9977 |

**Table 6:** $R^2$ of the Gaussian Fit of the First Three Expressions on the RHS of Equation 22 Plotted on Linear, Power Function-compressed, and Logarithmically Compressed Axes

| Linear | Power Function (0.5) | Power Function (0.33) | Logarithm |
|---|---|---|---|
| 0.5238 | 0.8536 | 0.904 | 0.9938 |
| 0.5339 | 0.8558 | 0.9035 | 0.9948 |
| 0.5374 | 0.8565 | 0.9034 | 0.9955 |

**Table 7:** $R^2$ for the Gaussian fit of the Expression 48 (with K=5 and N=1) for a few combinations of the Hill coefficients m and n

| | n=2 | n=4 | n=6 | n=8 | n=10 | n=12 |
|---|---|---|---|---|---|---|
| m=1 | 0.8442 | 0.9448 | 0.9872 | 0.9973 | 0.9952 | 0.9903 |
| m=2 | 0.8434 | 0.9377 | 0.9821 | 0.9965 | 0.9983 | 0.9947 |
| m=4 | 0.8275 | 0.9365 | 0.974 | 0.9916 | 0.9979 | 0.9979 |

The next two tables (8 & 9) show the average and standard deviation (in the parenthesis) of the slope, the y-intercept, and $R^2$ of the straight line fits for the nine combinations of $\beta_2/\alpha_2$ and $\beta_3/\alpha_3$ for the series of FNM and C1-FFL, respectively. The Weber fraction is equal to the following values: 0.001, 0.02, 0.05, 0.1, 0.2, 0.5, and 1.0,

**Table 8:** The Stochastic Validation for the Series of FNM

| Weber fraction | Avg Slope of Mu | Avg y-intercept of Mu | Avg Slope of Sigma | Avg y-intercept of Sigma | Avg $R^2$ for Mu | Avg $R^2$ for Sigma |
|---|---|---|---|---|---|---|
| 0.01 | 1.45 (3.4E-06) | 7.50E-04 (1.8E-06) | 1.39 (4.4E-04) | 3.70E-06 (4.5E-06) | 1 (0) | 1 (0) |
| 0.02 | 1.45 (1.9E-06) | 7.32E-04 (4.5E-06) | 1.39 (6.9E-04) | 1.84E-05 (1.5E-05) | 1 (0) | 1 (0) |
| 0.05 | 1.45 (3.5E-05) | 7.51E-04 (4.4E-05) | 1.39 (1.25-03) | -3.31E-05 (6.5E-06) | 1 (0) | 1 (0) |
| 0.1 | 1.45 (4.2E-05) | 6.83E-04 (6.0E-05) | 1.39 (7.2E-04) | 1.76E-04 (9.2E-05) | 1 (0) | 1 (0) |
| 0.2 | 1.45 (1.8E-05) | 8.41E-04 (5.1E-06) | 1.39 (1.2E-04) | 3.84E-04 (3.8E-05) | 1 (0) | 1 (0) |
| 0.5 | 1.80 (7.2E-02) | -4.40E-02 (2.1E-03) | 2.12 (1.2E-01) | -3.0E-02 (1.5E-03) | 9.99E-01 (9.6E-05) | 9.99E-01 (1.9E-04) |
| 1 | 5.70 (7.9E-01) | -9.3E-02 (1.9E-01) | 8.59 (1.4) | -2.40E-01 (3.1E-01) | 9.96E-01 (8.8E-04) | 9.96E-01 (1.0E-03) |

The value in the parenthesis is the standard deviation.

**Table 9:** The Stochastic Validation for the Series of C1-FFL

| Weber fraction | Avg Slope of Mu | Avg y-intercept of Mu | Avg Slope of Sigma | Avg y-intercept of Sigma | Avg $R^2$ for Mu | Avg $R^2$ for Sigma |
|---|---|---|---|---|---|---|
| 0.01 | 9.00E-01 (3.3E-07) | 6.77E-03 (4.7E-07) | 1.26 (1.0E-04) | -5.96E-04 (8.6E-07) | 1 (0) | 9.99E-01 (1.5E-07) |
| 0.02 | 9.00E-01 (1.3E-05) | 6.77E-03 (1.8E-06) | 1.26 (1.4E-04) | -1.18E-03 (2.0E-06) | 1 (0) | 9.99E-01 (1.5E-07) |
| 0.05 | 9.00E-01 (1.6E-05) | 6.76E-03 (2.2E-06) | 1.26 (4.6E-05) | -2.98E-03 (3.2E-06) | 1 (0) | 9.99E-01 (1.8E-08) |
| 0.1 | 9.00E-01 (1.1E-04) | 6.71E-03 (7.5E-05) | 1.26 (1.1E-04) | -5.95E-03 (2.3E-05) | 1 (0) | 9.99E-01 (3.8E-07) |
| 0.2 | 9.00E-01 (5.4E-05) | 6.51E-03 (8.2E-06) | 1.26 (6.5E-04) | -1.20E-02 (2.2E-04) | 1 (0) | 9.99E-01 (1.6E-06) |
| 0.5 | 9.04E-01 (4.2E-03) | 2.22E-01 (4.0E-02) | 1.30 (1.7E-02) | 2.49E-01 (4.5E-02) | 9.92E-01 (2.3E-03) | 9.80E-01 (6.3E-03) |
| 1 | 8.77E-01 (2.1E-03) | 3.84E-01 (3.0E-02) | 1.23 (9.9E-03) | 8.16E-01 (9.2E-02) | 9.98E-01 (9.9E-01) | 9.98E-01 (2.1E-03) |

The value in the parenthesis is the standard deviation.
